# Dynamic landscape and genetic regulation of RNA editing in schizophrenia

**DOI:** 10.1101/485086

**Authors:** Michael S. Breen, Amanda Dobbyn, Qin Li, Panos Roussos, Gabriel E. Hoffman, Eli Stahl, Andrew Chess, Pamela Sklar, Jin Billy Li, Bernie Devlin, Joseph D. Buxbaum, for the CommonMind Consortium (CMC)

## Abstract

RNA editing is vital for neurodevelopment and the maintenance of normal neuronal function. We surveyed the global landscape and genetic regulation of RNA editing across several hundred schizophrenia and control postmortem brain samples from the CommonMind Consortium covering two regions, the dorsolateral prefrontal cortex (DLPFC) and anterior cingulate cortex. In schizophrenia, RNA editing sites encoding AMPA glutamate receptors and post-synaptic density genes were less edited, while more editing was detected in sites implicated in translational initiation. These sites replicate between brain regions, map to 3’UTRs, enrich for common sequence motifs and coincide for RNA binding proteins crucial for neurodevelopment. Importantly, these findings cross-validate in hundreds of non-overlapping DLPFC samples. Furthermore, ~30% of RNA editing sites associate with cis-regulatory variants (edQTLs). Fine-mapping edQTLs with schizophrenia GWAS loci revealed colocalization of 11 edQTLs with 6 GWAS loci. This supports a causal role of RNA editing in risk for schizophrenia. Our findings illustrate widespread altered RNA editing in schizophrenia and its genetic regulation, and shed light onto RNA editing-mediated mechanisms in schizophrenia neuropathology.

## INTRODUCTION

Schizophrenia (SCZ) is a severe psychiatric disorder affecting ca. 0.7% of adults that is characterized by abnormalities in thought and cognition^1^. While the onset of SCZ typically does not occur until late adolescence or early adulthood, there is strong support from clinical and epidemiological studies that SCZ reflects a disturbance of neurodevelopment^2^. There is clear and consistent evidence that SCZ is largely a genetic disorder. Large-scale mapping of genetic risk variants has identified multiple rare copy number variants^3^, several rare single nucleotide variants^4,5^, and >100 common genetic loci^6^, the latter exerting small polygenetic effects on disease risk. This observation of a highly polygenic architecture has been widely replicated^7,8^. However, the role of sequence variation arising as a result of post-transcriptional events, such as RNA editing, remains largely unexplored.

RNA editing is a modification of double-stranded pre-mRNA that can introduce codon changes in mRNA through insertions, deletions or substitutions of nucleotides and hence can lead to alterations in protein function. Adenosine to inosine (A-to-I) editing is the most common form of RNA editing, affecting the majority of human genes and is highly prevalent in the brain^9,10^. These base-specific changes to RNA result from site-specific deamination of nucleotides catalyzed by adenosine deaminases acting on RNA (ADAR) enzymes, whereby a genetically encoded adenosine is edited into an inosine, which is read by the cellular machinery as a guanosine. Editing sites in coding regions can be conserved across species and are commonly located in genes involved in neuronal function^11,12^. RNA editing has been reported to modulate excitatory responses, permeability of ion channels and other neuronal signaling functions^13,14^. These sites have been shown to be tightly and dynamically regulated throughout pre- and post-natal human cortical development^15^. Aberrant RNA editing has also been reported in several neurological disorders, including major depression^16^, Alzheimer’s disease^17^, and amyotrophic lateral sclerosis^18^.

In SCZ, the role of RNA editing in serotonin and glutamate receptors has drawn significant attention largely due to the serotonergic and glutamatergic hypotheses of mood disorders. To this end, RNA editing research in SCZ has so far focused on targeted approaches in serotonin 2C receptor (5-HT_2c_R) and two classes of ionotropic glutamate receptors, 2-amino-3-(3-hydroxy-5-methyl-isoxazol-4-yl)-propanoic acid (AMPA) and kainate receptors. It has been shown that 5-HT_2C_R pre-mRNA undergoes editing at five sites on exon V, which results in amino acid changes and can produce >20 different receptor isoforms, each with specific activity^20,21^. The extent of editing in 5-HT_2C_R correlates with functional activity of the receptor and this has been reported in post-mortem brain tissue of individuals with major depression^16^ and schizophrenia^21,22^. Regarding glutamate receptors, AMPA and kainate receptor pre-mRNA undergoes editing at two sites (Q/R and R/G), both of which have functional significance^9,12^. The Q/R site occurs at a stably high editing rate with 100% editing, whereby loss of editing at this site causes enhanced Ca^2+^ permeability, possibly resulting in cellular dysfunction in SCZ^12,23,24^. The R/G site is not fully edited, which changes the kinetics of desensitization^25^. However, there are still a limited number of studies measuring RNA editing levels of these receptors in SCZ, and those which have been conducted report on relatively small sample sizes and often lack an unbiased genome-wide approach. Consequently, there is no consensus on the type of editing nor how pervasive altered RNA editing is in the brain of SCZ patients. Moreover, the underlying *cis*-acting genetic variants, which are associated with RNA editing levels (edQTLs) in the brain and whether these variants are also implicated in disease risk also remain poorly understood. To address these questions, integrative genome-wide studies of large patient cohorts and multiple implicated brain regions are needed.

The primary goal of the current investigation was to clarify the relevance of RNA editing in pathophysiology of SCZ using an unbiased, genome-wide approach applied to a large cohort of SCZ cases and control samples generated from the CommonMind Consortium (CMC), which is orders of magnitude larger than prior RNA editing studies. Two brain regions implicated in neurodevelopment and SCZ neuropathology were examined, including the dorsolateral prefrontal cortex (DLPFC; Brodmann areas 9 and 46) and anterior cingulate cortex (ACC). Results from this cohort were then reproduced in a separate, non-overlapping DLPFC cohort generated through the National Institute of Mental Health (NIMH) Human Brain Collection Core (HBCC). By applying a multi-step analytic framework and including genome-wide characterization of common genetic variation (**Figure S1**), we generated a resource of the genetics of RNA editing in the brain. We use this resource to identify: (1) genes and RNA editing sites with significant differences in RNA editing levels between subjects with SCZ and control subjects; (2) coordinated editing (co-editing) of RNA editing sites implicated in SCZ; and (3) specific effects on RNA editing of genetic variants previously implicated in disease risk. Our results shed light on the subtle effects expected from the polygenic nature of SCZ risk and thus substantially refine our understanding of the RNA editing mediated mechanism involved in the neurobiology of SCZ.

## RESULTS

### Discovery and validation samples

In order to quantify RNA editing events, we leveraged RNA-sequencing data from post-mortem brain tissue collected and generated on behalf of the CMC. Two brain regions, including the ACC (SCZ=225, Controls=245) and the DLPFC (SCZ=254, Controls=286) were investigated, and together these samples served as the *discovery cohort* (**Figure S2**). These samples were also genotyped on the Illumina Infinium HumanOmniExpressExome array. In parallel, we also leveraged a completely separate, non-overlapping cohort consisting of post-mortem DLPFC tissue (SCZ=100, Controls=204) collected and generated on behalf of NIMH HBCC. This second resource served as a *validation cohort* so as to cross-validate SCZ-related editing events identified from the CMC discovery samples.

### Overall RNA editing levels in SCZ

We first asked whether overall levels of RNA editing varied between SCZ and control samples, separately in the ACC and DLPFC. Overall editing levels were computed within each individual sample defined as the percentage of edited nucleotides at all known editing sites. We did not impose any coverage criteria, but instead took all sites into account that were used in this study to obtain the total amount of editing in each sample. Higher levels of overall RNA editing in SCZ cases were observed compared to controls in the ACC and DLFPC (*p*=0.0001, *p*=7.2×10^−6^, respectively) (Figure 1A). Approximately 30% of the variation in overall RNA editing levels was explained by ADAR1 (*p*=<2.2×10^−16^), ADAR2 expression explained 17% of variation in overall RNA editing (*p*=1.2×10^−08^) and ADAR3 expression had no significant effect (*p*=0.10) (Figure 1B-C). In addition, marked increases in overall editing levels were observed within definite genic regions, specifically 3’UTR and intergenic regions in SCZ compared to control samples, which replicated across the ACC and DLPFC (**Figure S3A**). Moreover, as previous research has quantified RNA editing levels explicitly in serotonergic and glutamatergic receptors, we also computed overall editing levels in serotonin and glutamate receptor activity genes using *a priori* defined gene-sets (GO:009589 and GO:0008066, respectively). To this end, higher levels of overall editing were found in glutamate receptors in SCZ cases relative to controls in the ACC (*p*=0.0004) and DLPFC (*p*=0.001), while no significant differences were found in the levels of overall editing in serotonin receptors (**Figure S3B**). Expression of ADAR1 and ADAR2 were also significantly higher in SCZ compared to control samples (**Figure S3C-E**). Importantly, these observations were collectively and robustly reproduced in our independent, non-overlapping DLPFC validation cohort, which together highlight higher overall levels of RNA editing in SCZ, especially within 3’UTR and intergenic regions, as well as within genes encoding glutamatergic receptors.

**Figure 1.**
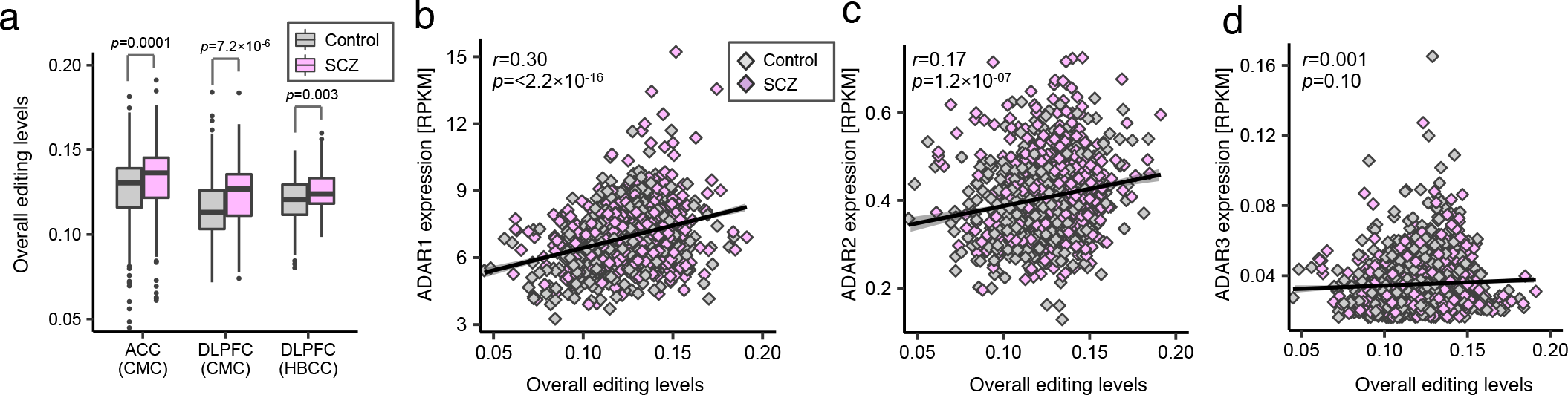
Overall RNA editing profiles. (**a**) Overall RNA editing levels across all known RNA editing events for were computed for each individual and subsequently compared between SCZ and control samples. We define overall RNA editing as the total number of edited reads at all known RNA editing sites over the total number reads covering all sites. A Mann-Whitney U test was used to test significance between groups. Associations between expression levels of (**b**) ADAR1, (**c**) ADAR2 and (**d**) ADAR3 (quantified as the number of RNA-seq reads per kilobase of transcript per million mapped reads (RPKM)) and overall editing levels across all available ACC and DLPFC samples (including CMC and HBCC data). R-values were calculated by robust linear regressions on overall editing levels and logarithmic transformed RPKM values.

To rule out the possibility that these reproducible differences in overall RNA editing levels may be driven by medication effects, we examined overall editing levels in postmortem DLPFC tissue derived from an RNA-sequencing study of 34 Rhesus macaque monkeys treated with high doses of haloperidol (10 mg/kg/d), low doses of haloperidol (4mg/kg/d), clozapine (5.2 mg/kg/d), and vehicle. We found no associations between overall RNA editing levels with medication or dosage (**Figure S4**), indicating that antipsychotic treatments likely do not have a strong effect on the amount of overall RNA editing observed in SCZ cases.

### Discovery of altered RNA editing sites in SCZ

We next set out to identify RNA editing sites associated with SCZ by testing if the degree of RNA editing levels at each site is significantly different between SCZ and control samples. To this aim, a compendium of high quality and high confident RNA editing sites was assembled by imposing a series of detection-based thresholds. Known single nucleotide variants were removed from subsequent analyses and stringent filters for base quality, mapping quality and coverage were used (see Methods). Next, we required RNA editing events to be detected in at least 70% of all samples. After these quality control and filtering steps, we identified a high-confidence set of 11,242 RNA editing sites in the ACC and 7,594 sites in the DLPFC, for which there were no systematic differences in the mapping, base quality, and read coverage between SCZ and control samples. A significant fraction of these RNA editing events replicated across brain regions (∩^sites^=6,999, OR=21.05, *p*=<2.0×10^−50^). A large fraction of these sites were located in Alu repeat elements, mapped to 3’UTR regions and were enriched for A-to-I conversions (**Figure S5**).

Following this curation of editing sites, differential RNA editing analysis was carried out. It is likely that genome-wide RNA editing events, similar to gene expression profiles, may be influenced by differences in biological and technical factors. To this end, a linear mixed effect model was applied to decompose the computed RNA editome into the percentage attributable to multiple biological and technical sources of variation. For each RNA editing site, the percentage of editing variation was computed attributable to individual’s age, RIN, PMI, pH, brain weight, library batch, ethnicity, gender, medication and various sample sites. These variables displayed little influence on RNA editing profiles, with individual age having the largest genome-wide effect and explained a median 0.79% of the observed variability (**Figure S6A**). These factors, however, explained a much higher amount of median variability in matching gene expression profiles than observed RNA editing profiles (**Figure S6B**). Subsequently, differential editing analysis covarying for individual’s age, RIN, PMI, sample site and sex identified 182 sites in the ACC and 194 sites in the DLPFC significantly associated with SCZ (Adj. *P*-value < 0.05) (Figure 2A; **Table S1A-B**). Among the top-ranked sites, were those encoding for genes *ATRLN1*, *AKAP5* and *RPS20* in the ACC and *KCNIP4*, *VPS41* and *ZNF140* in the DLPFC. A high degree of concordance was observed between the altered RNA editing sites in the ACC and DLPFC (*R^2^*=0.77) and a significant overlap of differentially edited sites replicated between brain regions (∩^sites^=29, OR=12.6 *p*=9.6×10^−20^) (Figure 2B). Differentially edited sites were also enriched in genes found to be highly expressed in the ACC (∩^sites^=42, OR=5.4 *p*=2.0×10^−13^) and DLPFC (∩^sites^=54, OR=8.6 *p*=2.7×10^−22^) and that the majority of genes with differential RNA editing sites did not display differential gene expression (**Figure S7A-F**). Moderate, yet significant, correlations were also observed between RNA editing levels and gene expression, implying RNA editing as a possible posttranscriptional mechanism for the regulation of gene expression (**Figure S7G-I**).

**Figure 2.**
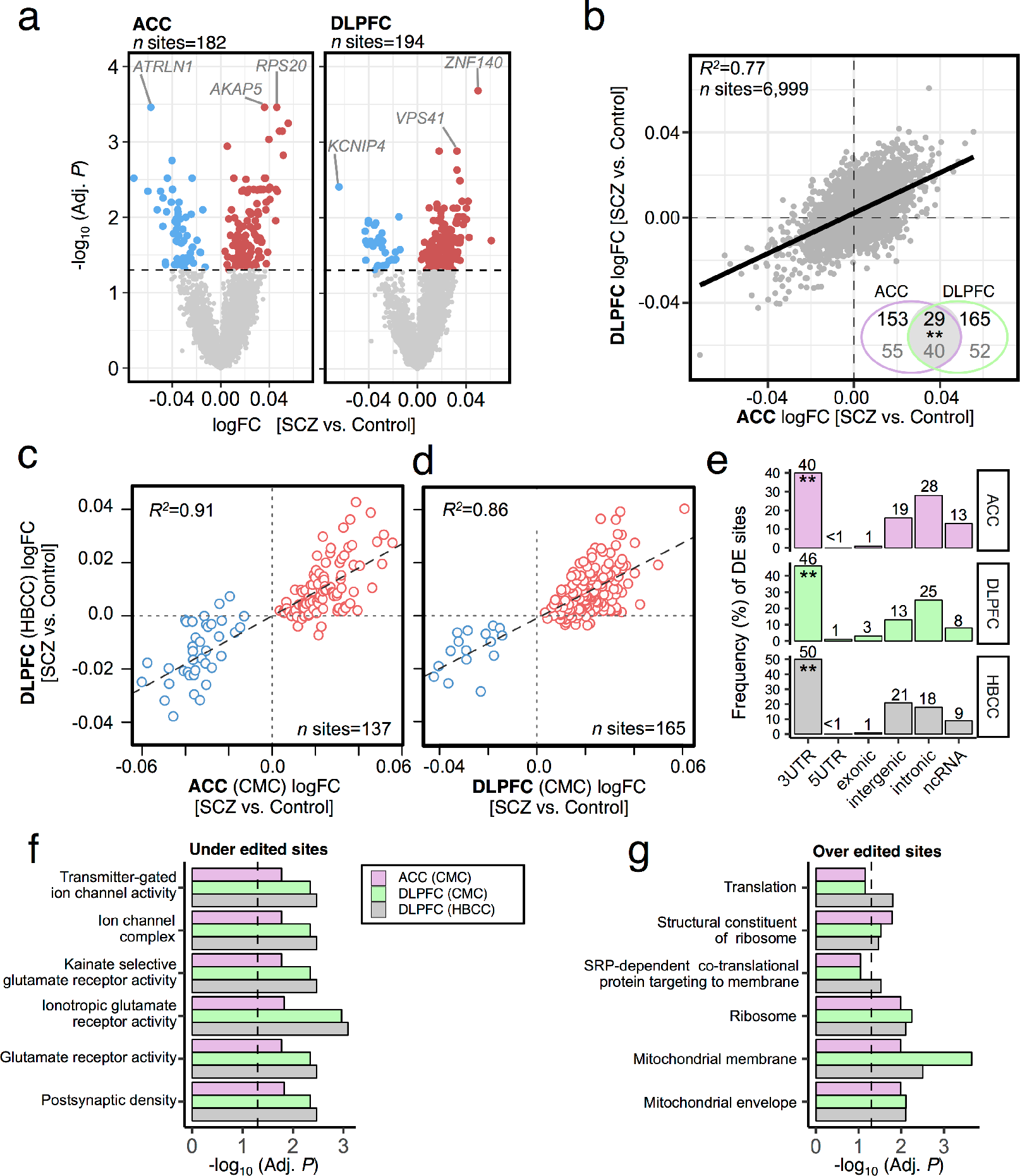
Differential RNA editing analysis. Identification of differentially edited sites in SCZ cases (**a**) within the ACC and DLPFC. The dotted line marks a multiple test corrected level of significance (Adj. P < 0.05). Red dots indicate over-edited sites and blue dots indicate under-edited sites. For the top three sites, we outline their respective gene body. (**b**) Overall concordance between the extent of SCZ-related log fold changes of RNA editing sites in the ACC versus DLPFC. The inset Venn Diagram indicates the total number of genome-wide significant sites (top value) and respective gene symbols (bottom value), which overlap between brain regions. Scatterplots of log fold changes for (**c**) 137 differentially edited sites in the ACC (x-axis) and (**d**) 165 differentially edited sites in the DLPFC (x-axis), respective to log fold change within independent HBCC DLPFC samples (y-axis). (**e**) Significantly differentially edited sites by genic region indicates a significant enrichment (P < 0.05,**) of sites mapping to 3’UTR regions. Enrichment was calculated using a Fisher’s exact test with background set to all 3’UTR sites within the RADAR database. Functional annotation of the top five enrichment terms for (**f**) under-edited sites and (**g**) over-edited sites in SCZ. **Data information:** R^2^ values were calculated by robust linear regressions on log fold change differences.

### Validation of altered RNA editing sites in SCZ

Next, we asked whether these differential editing patterns in SCZ replicate within our independent, non-overlapping DLPFC validation cohort. These samples underwent matching quality-control metrics to identify a collection of high confidence RNA editing events, as noted above. A total of 15,0 RNA editing events were detected across these validation samples and a significant fraction of sites were also detected in the ACC (∩^sites^=8354, OR=4.01 *p*=5.6×10^−251^) and DLPFC (∩^sites^=6659, OR=7.73 *p*=4.67×10^−248^) discovery samples (**Table S2**). Differential RNA editing analysis was carried out on these independent samples as previously described. In order to assess replication, we first measured the concordance between directionality of change statistics (log fold-change) for the differentially edited sites identified in the ACC and the DLPFC discovery samples relative to these independent DLPFC validation samples. High levels of concordance were observed for the candidate SCZ-related RNA editing sites in both the ACC and DLPFC (*R^2^*=0.91, *R^2^*=0.86, respectively) (Figure 2C-D). Subsequently, two prediction models were built based on differentially edited sites from the (1) DLPFC and (2) ACC discovery samples using regularized regression models and evaluated their performance to predict class labels (*i.e.* distinguish between SCZ and control samples) on withheld DLPFC validation samples. Classification accuracies were reported as area under the receiver operative curve on withheld DLPFC samples. When distinguishing between SCZ and control samples, classification accuracies reach 78% and 72% on withheld, independent DLPFC samples when using differentially edited sites derived from DLPFC and ACC discovery samples, respectively (ridge regression outperformed other methods: **Figure S8**). Overall, these results suggest a moderate level of cross-validation of SCZ-related editing events across brain regions and independent cohorts.

### Characterization of differentially edited sites

Differentially edited sites derived from discovery (ACC and DLPFC) and validation (DLPFC) samples were comprehensively annotated. A significant fraction of differentially edited sites consistently mapped to 3’UTRs across brain regions and cohorts (Figure 2E). Functional enrichment analysis revealed that under-edited sites consistently mapped to postsynaptic density genes as well as genes encoding kainate and glutamate receptor activity and over-edited sites mapped to genes implicated in protein translation and mitochondrial-related terms (Figure 2F-G). We also examined whether these differentially edited sites map to genes with specific developmental expression profiles using gene expression data from the BrainSpan Project and found that differentially edited sites in SCZ consistently mapped to genes that are predominately postnatally biased in expression (**Figure S9**). These genes were found to peak in brain expression during young and middle adulthood, developmental windows when SCZ often becomes clinically recognizable.

As these sites share several sequence and functional features, we explored whether differential editing sites may share a common sequence motif potentially important for editome recognition (20±nt centered on targeted A nucleotide). Consistent and strong enrichment was found for a 10-nt motif (CTGGG<u>A</u>TTACA) in region adjacent to most differential editing sites located in 3’UTR regions (**Figure S10, Table S3**). This short sequence has been reported to occur frequently within noncoding regions and is also found to overlap with fragments of Alu repeat elements. Subsequently, we examined whether differentially edited sites found to share this sequence motif also mapped to any known human RNA binding protein (RBP) binding sites (30±nt centered on target A). A large fraction of these sites (>61% in each brain region) significantly coincided with binding sites specific for RBP serine/arginine (SR)-rich splicing factor 5 (*SRSF5*) (**Table S4**). Interestingly, this protein is associated with pyruvate carboxylase deficiency, a disorder that leads to developmental delay, recurrent seizures and a failure to produce the necessary the enzymes that facilitate the production of neurotransmitters and meet the energy demands required for typical brain development. Other significant RBP binding sites included additional members of the SR-rich family of pre-mRNA splicing factors, such as *SRSF2* and *SRSF3* as well as CUG triplet repeat binding protein (*CUGBP*), an RBP found to mediate neuronal toxicity. Taken together, these results indicate that differentially edited sites replicate across brain regions and cohorts, implicate altered glutamatergic receptor activity and protein translation, further impact 3’UTR regions and are likely edited by a similar editing mechanism (and possible RBPs) as indicated from a commonly identified sequence motif.

### Genes enriched with differential RNA editing in SCZ

We examined whether any genes contained an enrichment of differentially edited sites beyond what could be expected by chance. As expected, gene length functions as a correlate of the total number of RNA editing sites per gene (**Figure S11**). Therefore, we computed over-representation of differential RNA editing sites within each gene by setting a rotating background specific to the total number of known RNA editing events for a particular gene in order to systematically correct for gene length (**Table S5**). Genes enriched for under-edited sites in SCZ were primarily located within intronic regions while genes enriched for over-edited sites were primarily located in 3’UTRs (Figure 3A). Three genes, including *KCNIP4*, *HOOK3* and *MRPS16* displayed enrichment for altered editing sites across the ACC, DLPFC and our independent validation DLPFC cohort (Figure 3B,D,E). *KCNIP4* harbored 13 unique differentially edited sites spread over its first and second introns, which were predominately under-edited in SCZ compared to control samples. *KCNIP4* is a member of the voltage-gated potassium channel-interacting proteins and has been shown to interact with presenilins and modulate pacemaker neurons in the reward circuitry of the brain^26,27^. Genome-wide association studies (GWAS) have also found *KCNIP4* to be associated with SCZ, suicidal ideation and attention-deficit/hyperactivity disorder^28–30^. *HOOK3* harbored 22 unique sites and *MRPS16* harbored 19 unique sites both within their respective 3’UTR regions, which we repredominately over edited in SCZ compared to control samples. *HOOK3* is a microtubule tethering protein essential for centrosomal assembly during neurogenesis and brain development^31^ and *MRPS16* is a mitochondrial ribosomal protein involved in mitochondrial protein translation^32^.

**Figure 3.**
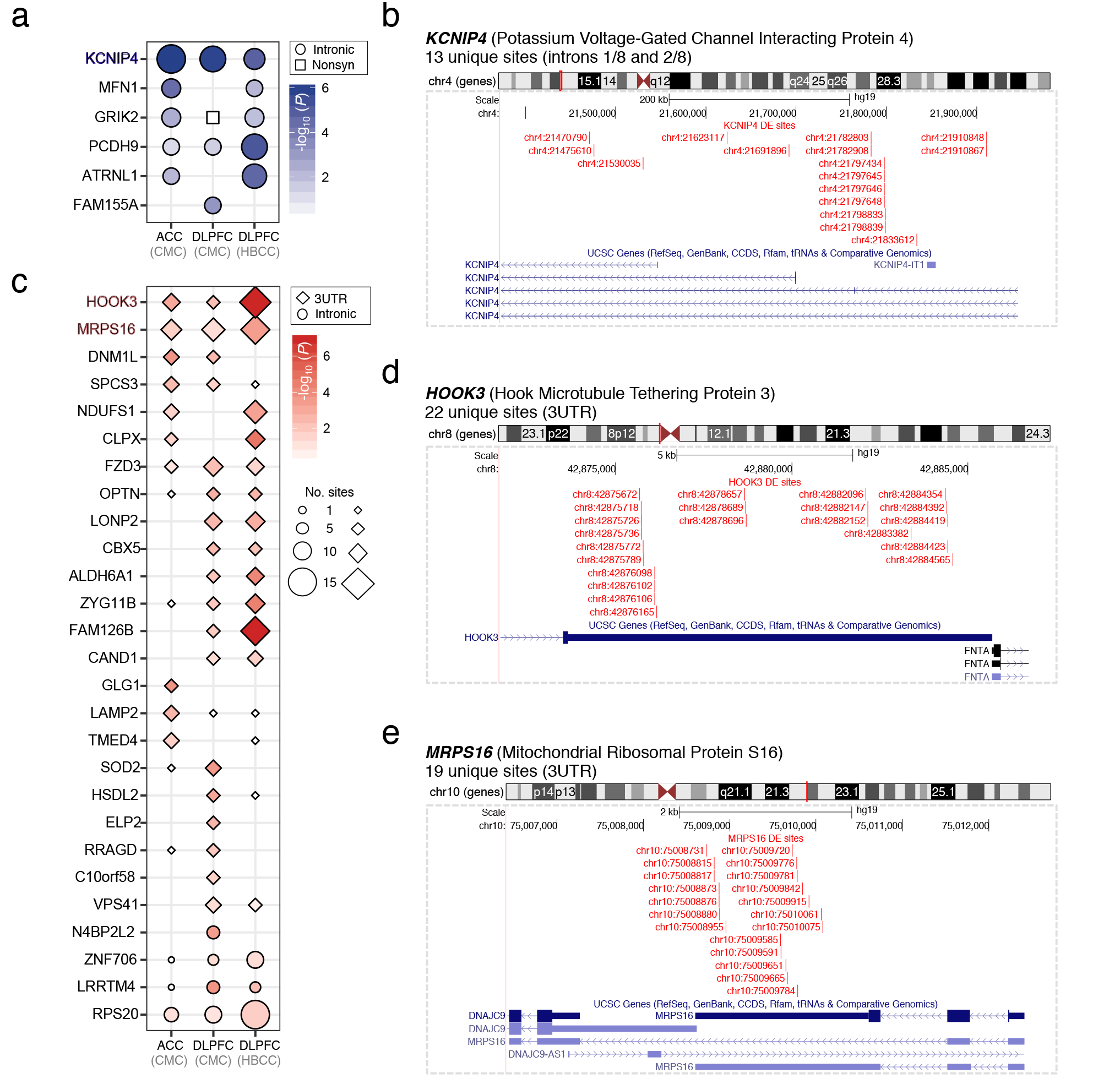
Genes enriched with differential editing. (**a**) Genes enriched for under-edited sites in SCZ primarily map to intronic regions. (**b**) KCNIP4 contains 16 unique differential RNA editing sites, which are under-edited and span its first and second intron. These sites replicate across brain regions and withheld validation samples. (**c**) Genes enriched for over-edited sites in SCZ primarily map to 3’UTR regions. (**d**) HOOK3 contains 22 unique differential RNA editing sites and (**e**) MRPS16 contains 19 unique differential RNA editing sites which are over-edited within their respective 3’UTR region. **Data information:** UCSC Genome Browser customized track options were used to display the precise locations of editing sites within each gene.

### Co-editing networks associate with SCZ

Discrete groups of coordinately edited (co-edited) sites were identified and tested for association to SCZ using an unbiased network approach, separately for ACC and DLPFC discovery samples. A total of five co-editing modules were detected in each brain region and displayed a near one-to-one mapping between the ACC and DLPFC (Figure 4A), indicating highly similar co-editing network topology. Modules were assessed for over-representation of differential RNA editing sites and two modules were identified in the ACC (M1a and M4a) and two modules in the DLPFC (M1d and M4d) (Figure 4B). Module eigengene (ME) values for these modules elucidated higher levels of editing in modules M1a and M1d and lower levels of editing in modules M4a and M4d in SCZ compared to control subjects (Figure 4C). Functional annotation of over-edited modules M1a and M1d revealed strong enrichment for regulation of translation and translation initiation, while under edited modules M4a and M4d were enriched for AMPA glutamate and ionotropic receptors (Figure 4D). Cell type enrichment analysis revealed modules M1a and M1d were enriched for pyramidal neurons while modules M4a and M4d were enriched for interneurons (Figure 4E). Moreover, M1a and M1d were positively associated with ADAR1 and ADAR2 expression and modules M4a and M4d were negatively associated with ADAR2 expression (**Figure S12**). Upon closer inspection, several sites located within modules M4a and M4d mapped to nonsynonymous sites in genes *NOVA1*, *UNC80*, *GRIA2*, *GRIA3*, *GRIA4*, *GRIK2* and *ANKD36*, and these sites were predominately under-edited in SCZ compared to control samples (Figure 4F). Several of these sites, particularly the Q/R and R/G sites in *GRIA2*, are well documented as fully edited sites under normal conditions whereby loss of editing in these sites leads to enhanced Ca^2+^ permeability and cellular dysfunction, and this has been suggested to play a role in SCZ^23,24^. *NOVA1* is essential for normal postnatal motor function and regulates alternative splicing of multiple inhibitory synaptic targets^32^. *NOVA1* has been reported to be dysregulated at the gene level in independent SCZ postmortem brain samples^32^ and RNA editing in *NOVA1* has been shown to influence protein stability^33^, but has yet to be associated with SCZ.

To validate these findings, a separate co-editing network was constructed for our DLPFC validation cohort. A total of six co-editing modules were detected, which displayed high degree of overlap with co-editing modules previously identified within our discovery cohort (**Figure S13A**). Two modules harbored a significant fraction of differentially edited sites (M1h and M4h) and ME values for M1h displayed higher levels of editing in SCZ whereas module M4h displayed lower levels of editing in SCZ compared to control samples (**Figure S13B-C**). Importantly, module M1h harbors a significant number of sites in common with modules M1a (∩^sites^=159, OR=46.77 *p*=2.0×10^−156^) and M1d (∩^sites^=193, OR=50.29 *p*=2.0×10^−192^), and module M4h shares a significant number of sites with modules M4a (∩^sites^=31, OR=4.91, *p*=1.56×10^−11^) and M4d (∩^sites^=7, OR=3.34, *p*=0.016). M1h was significantly enriched for mitochondrial protein translation-related terms, whereas functional annotation of M4h displayed enrichment for AMPA glutamate receptor activity (**Figure S13D-E**). Moreover, M1h and M4h were both positively associated with ADAR1 and ADAR2 expression. These findings indicate replicated co-editing networks in SCZ across brain regions and cohorts, which together converge on decreased AMPA co-editing networks and increased protein translation modules.

**Figure 4.**
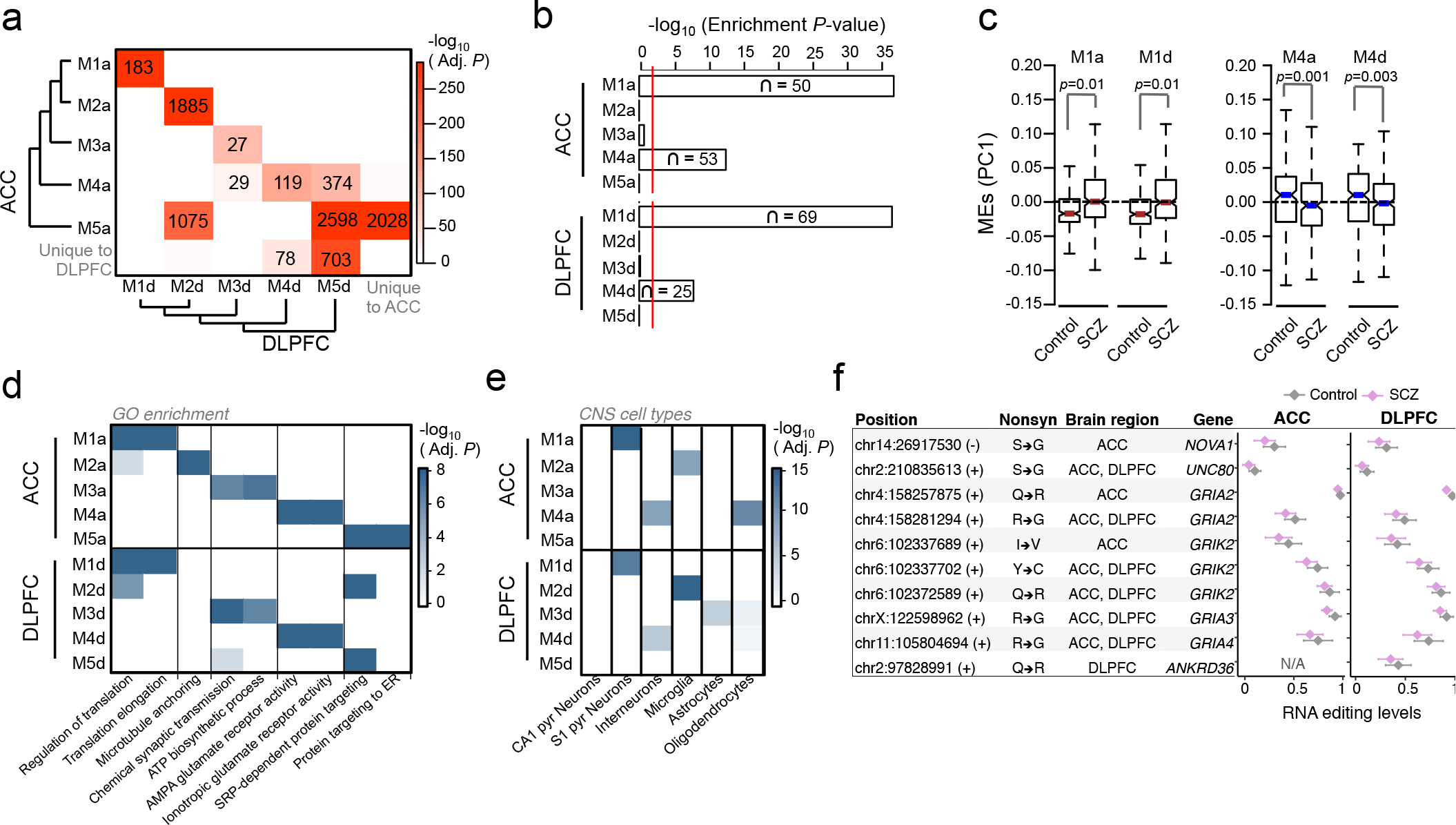
Unsupervised co-editing network analysis. (**a**) Overlap analysis of co-editing modules identified wi the ACC and DLPFC. Unsupervised clustering was used to group modules by module eigengene (ME) values using Pearson’s correlation coefficient and Ward’s distance method. Significance of overlap was computed using a Fis exact test corrected for multiple comparisons and colored on a continuous scale (bright red, strongly significant; wh no significance). The number of overlapping sites are displayed in each cell with a significant overlap, (**b**) Enrichm analysis of differentially edited sites within co-editing networks, (**c**) Assessment of ME values for modules M1a; M1d (over-edited) and M4a and M4d (under-edited). Differential ME analysis was conducted using a linear model < covarying for age, RIN, PMI, sample site and gender, (**d**) The top functional enrichment terms and (**e**) brain cell-type enrichment results for all identified modules, verifying similar functional and cell-type properties of co-editing netwc in the ACC and DLPFC. (**f**) A collection of nonsynonymous sites within SCZ-related AMPA glutamate recef. modules M4a and M4d (** indicates Adj. P <0.05, * indicates P < 0.05 derived from differential RNA editing analysis

### Identification and characterization of brain cis-edQTLs

Whole-genome genotype data were available for ACC and DLPFC samples used in our discovery cohort and were imputed using standard techniques, as previously described^7^. Genotype data were used to detect SNPs that have an effect on RNA editing levels (edQTL, editing quantitative trait loci). RNA editing levels from European-ancestry samples (ACC *N*=360; DLPFC *N*=421) were adjusted to fit a standard normal distribution and to reduce systematic sources of variation.

Adjusted editing levels were then fit to impute SNP genotypes, covarying for individual age, sample site and gender, PMI, RIN and diagnosis, using an additive linear model implemented in MatixEQTL. To identify genetic variants that could explain the variability of RNA editing, we first ran association tests between editing levels and genotypes by restricting the variant search space to only those within the same gene as each editing site and found an abundance of low *P*-values (**Figure S14**). Subsequently, we relaxed this assumption to define a broader window and identified 188,778 cis-edQTL (*i.e.* SNP-editing pairs ± 100kb of a site) in the ACC and 156,865 cis-edQTLs in the DLPFC at a genome-wide FDR < 5% (Figure 5A). Many of the edQTLs for the same site were highly correlated, due to linkage disequilibrium, and 70.9% of edQTL SNPs (edSNPs) in the ACC and 68.9% of edSNPs in the DLPFC predicted editing of more than one site. Notably, edQTLs tend to be present for editing sites with greater variance in editing levels (**Figure S15**). Each max-edQTL (defined as the most significant edSNP per site, if any) meeting a genome-wide significance threshold was located close to their associated editing site and acting in cis (5kb±nt) (Figure 5B, **Figure S15**). We reasoned that due to the propensity of edQTLs to be located close to their associated editing site, they should also influence additional editing sites nearby. This reasoning was strengthened by the observation that editing levels of editing sites within the same gene are more closely correlated than editing levels of editing sites in different genes (**Figure S16**).

**Figure 5.**
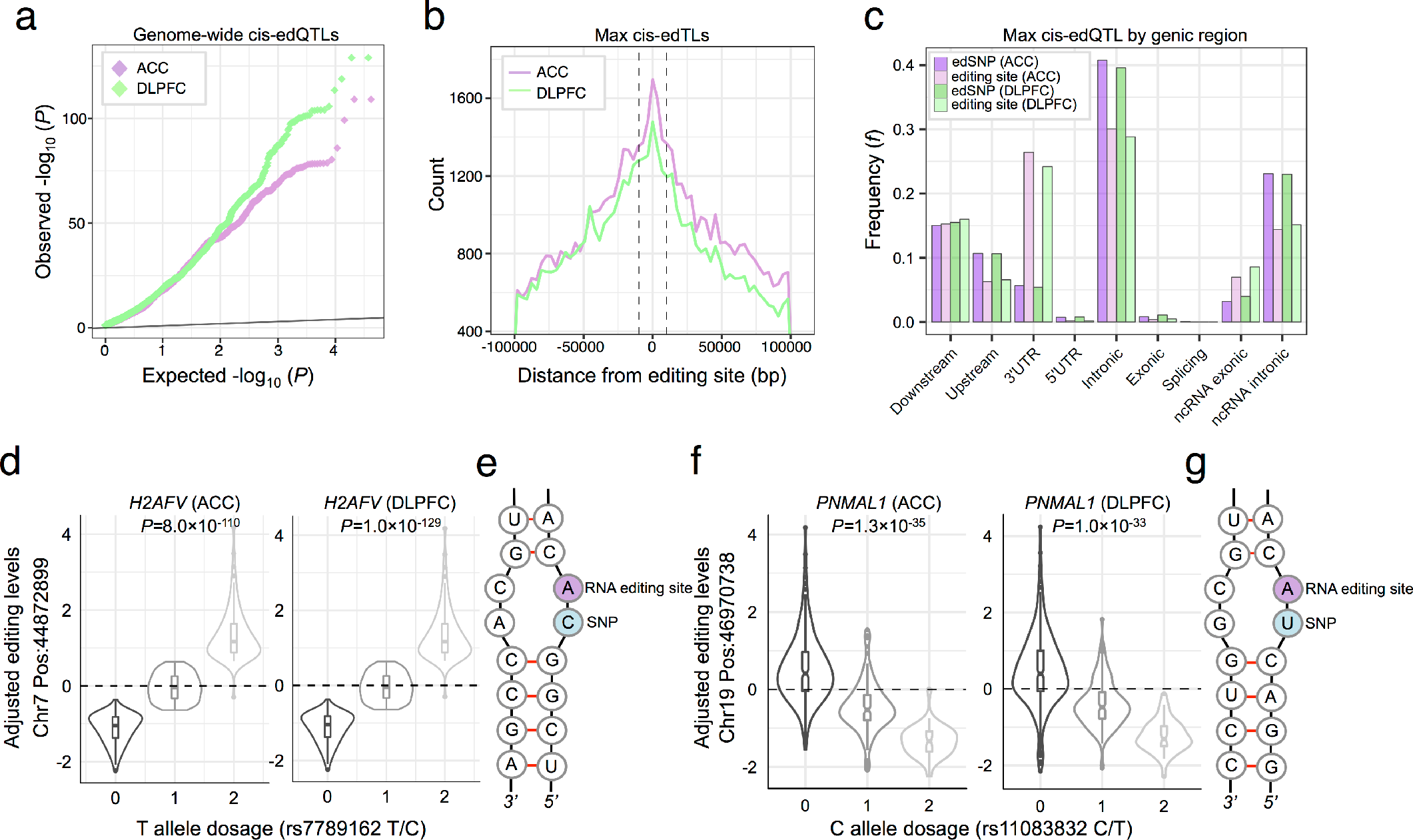
Brain cis-edQTL analysis. (**a**) Quantile-Quantile plot for association testing genome-wide P-values between genetic variants and 11,242 RNA editing sites in the ACC and 7,594 RNA editing sites in the DLPFC. (**b**) Distribution of the association tests in relation to the distance between the editing site and variant for max cis-edQTLs (that is, the most significant edSNP per site, if any). Vertical dotted lines indicate ± 5KB relative to the editing site. (**c**) Genic locations of edSNPs and corresponding editing sites. (**d-g**) Two examples of top cis-edQTLs with near by editing sites replicating between brain regions with (**e,g**) predicted local RNA secondary base-pairing structures.

Max-edQTLs in the ACC and DLPFC were enriched within genic elements and noncoding RNAs, particularly within intronic regions, while the corresponding editing sites were also enriched in 3’UTR regions (Figure 5C). Max-edQTLs edSNPs were also examined for tissue-specific enhancer specify using data from the FANTOM project across 40 different human tissues. edSNPs in the ACC and DLPFC were strongly enriched for brain-specific enhancer sequences more so than any other tissue (**Figure S17**). A significant fraction of max-edQTLs edSNPs replicated between the ACC (62%) and DLPFC (70%) (∩=34,367, Z-score=17,443, *p*=0.0009). Among the most significant associations identified in both brain regions were those in genes *H2AFV* and *PNMAL1*, where the edSNP is located immediately upstream of the RNA editing site within the same Alu (Figure 5D-G). In both cases, the alternative allele is unable to pair with the opposite base within the doublestranded RNA hairpin, introducing two consecutive mismatches in the local RNA secondary structure.

In addition, edSNPs were examined for association with gene expression levels by calculating the overlap between max-edQTLs and previously computed max expression QTL (max-eQTL) summary statistics derived from the ACC and DLPFC. A total of 29,335 edSNPs (54.4%) in the ACC were also associated with variation in gene expression, for which 31.3% were associated with a gene and one or more editing sites within the same gene (*e.g.* SNP_x_ is associated with Gene_y_ and one or more editing sites located within Gene_y_). Similarly, a total of 27,133 edSNPs in the DLPFC (55.6%) were also associated with gene expression variation, for which 30.1% were associated with a gene and one or more editing sites within the same gene.

### edQTL signatures co-localize with SCZ GWAS associations

It has previously been shown that a substantial proportion of schizophrenia GWAS associations (~20%) may be mediated by differential gene expression regulation^7^. RNA editing may represent an additional biological mechanism through which associated variants exert their effects on disease risk. Here, we leverage our edQTL resource to identify RNA editing sites that potentially alter SZC risk. Of the 108 SCZ GWAS loci reported previously, 14 harbor edQTL eSNPs for one or more RNA editing sites identified in either ACC or DLPFC. However, the presence of an edQTL within a GWAS locus does not imply disease causality. We therefore implemented coloc2, a Bayesian approach that integrates over statistics for all variants within a specified locus and estimates posterior probabilities of co-localization between two sets of association signatures, in order to identify RNA editing sites likely to contribute to SCZ etiology. We applied coloc2 to our ACC and DLPFC edQTL data in conjunction with summary statistics for the 108 genome-wide significant schizophrenia GWAS loci. We found evidence for co-localization (posterior probability > 0.5) of ACC edQTL and GWAS signatures at four loci comprising four unique edQTL and of DLPFC edQTL and GWAS signatures at four loci comprising seven unique edQTL (**Table S6**). Two of these loci are colocalized in both ACC and DLPFC; therefore a total of six GWAS associations, representing approximately five percent of all genome-wide significant loci, are potentially mediated by aberrant RNA editing (**Figure S18**). Of the six GWAS loci harboring SCZ-associated cis-edQTLs, these findings include genes, neuronal guanine nucleotide exchange factor (*NGEF*) and ADP-ribosylation-like factor 6 interacting protein 4 (*ARL6IP4*), which replicate between brain regions, as well as propionyl-CoA carboxylase subunit beta (*PCCB*) and long intergenic non-protein coding RNA 2551 (*RP11-890B15.3*), which are unique to the ACC and genes endosulfine alpha (*ENSA*) and diacylglycerol kinase iota (*DGKI*), which are unique to the DLPFC; one of which is highlighted in Figure 6.

**Figure 6.**
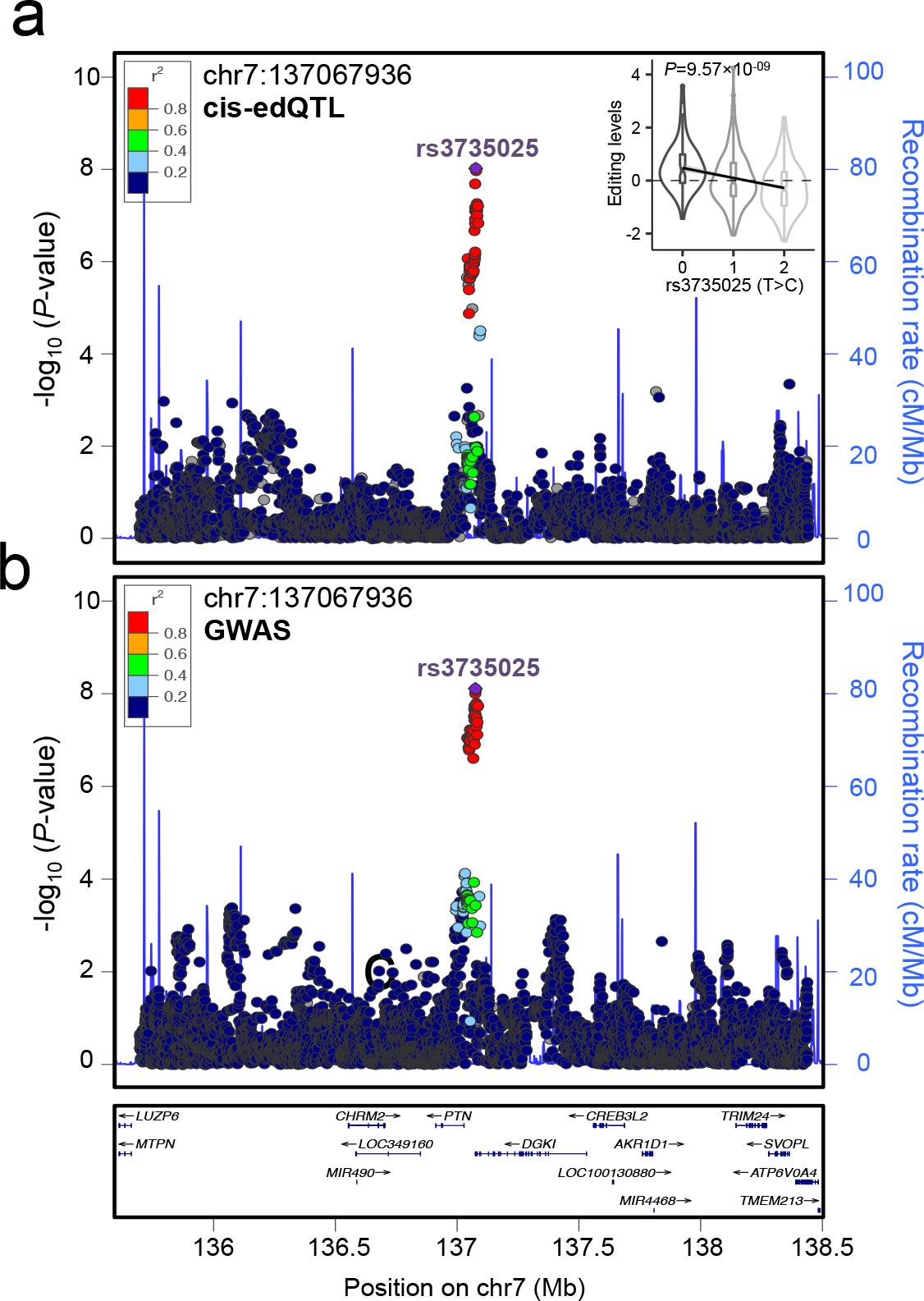
Coloc2 fine-mapping analysis. One example of co-localization between (**a**) cis-edQTL and (**b**) GWAS signal on chromosome 7 DGKI locus (PPH4=0.99). This specific co-localization event is specific to the DLPFC. LD estimates are colored with respect to the GWAS lead SNP (rs3735025, labeled) and coded as a heatmap from dark blue (0≥r^2^>0.2) to red (0.8≥r^2^>1.0). Recombination hotspots are indicated by the blue lines (recombination rate in cM Mb^−1^), (**a**) Inset violin plots reflects the association of editing between RNA editing site Chr7_127067936 (N = 421, P = 9.5 × 10^−09^) with SCZ risk allele at the GWAS index SNP in the respective loci.

## DISCUSSION

The recent expansion of RNA sequencing data sets has led to the identification of a huge number of RNA editing events, which affect the majority of human genes and is highly prevalent in the brain. Many such sites are commonly located in genes involved in neuronal maintenance and aberrant editing events have been associated with various neurological disorders. However, it has yet to be understood how pervasive RNA editing events are in the brain of SCZ patients and what are genetic forces guiding the regulation of these events. The ACC and DLPFC have been shown to play an important role in neurodevelopment and have been implicated in the pathophysiology of SCZ through abnormal regulation of executive function, social cognition, emotion, and self-reference. Here we have used genome-wide RNA-sequencing data derived from these tissue to advance our understanding of the function and regulation of A-to-I editing and to clarify how RNA editing and genetic liability is related to the molecular etiology of SCZ. We detected reproducible differences in RNA editing in SCZ that point to disruption of synaptic transmission, glutamatergic receptors and mitochondrial protein translation. Notably, these sites shared several common feature characteristics, including enrichment in 3’UTR regions, sharing similar sequence motifs and related patterns of co-editing in SCZ. We also identified several genes, including *KCNIP4*, *HOOK3* and *MRPS16*, which contained a large number of differentially edited sites in SCZ. Finally, by intersecting RNA-editing and genetics, we elucidated important aspects of the genetic control of editing and also found that genetic variants in 6 of the 108 SCZ GWAS risk loci alter editing of one or more editing sites.

The majority of differentially edited sites mapped to 3’UTR regions. As the vast majority of these sites are A-to-I conversions and are located in Alu elements, the stability of the resulting RNA structure is likely to be reduced^34^. Similarly, it is likely that RNA editing is contributing to the modulation of the stability of the folded pre-mRNA secondary structure in SCZ. To this end, lower levels of RNA editing were associated with postsynaptic density and glutamatergic genes as well as kainate and glutamate receptor activity genes (Fig. 2), likely resulting in lower stability in the double-stranded region after editing at these sites. Importantly, these genes comprise some of the most prominent and well-published genes in SCZ biology, including *GRIA2*, *GRIA3*, *GRIK1*, *GRIK2*, for which aberrant RNA editing levels have been well documented^23–25,35^, as well as *NRXN1* and *KALRN*, which have been less studied. Copy number variations and exonic deletions affecting the *NRXN1* gene have been associated with several neurodevelopmental disorders, including autism spectrum disorder^36^, SCZ^37^ and non-syndromic intellectual disability^38^, but its RNA mediated mechanisms are less understood. *NRXN1* generates multiple splice variants of the longer *α*-neurexin and shorter *β*-neurexin proteins, all of which function in synaptic adhesion, differentiation, and maturation^39^. *KALRN* is known to regulate neurite initiation, axonal growth, dendritic morphogenesis, and spine morphogenesis and is a key factor responsible for reduced densities of dendritic spines on pyramidal neurons in the DLPFC^40^, as previously reported in SCZ^41^. We also found enrichment for additional postsynaptic density and ion channel complex genes, including *RTN4*, *KCNIP4*, *NTNAP2* and *LRRTM4*. Notably, *KCNIP4* was enriched for differentially RNA editing sites in SCZ, which were found predominately spanning its first intron (Fig. 3). A major function attributed to *KCNIP4* is the regulation of the potassium channel Kv4, which are significant contributors to action potential activity in neurons. The first intron of *KCNIP4* is involved in alternative splicing events leading to Var IV of *KCNIP4*, which has been found to disrupt this current through failure to properly interact with presenilins, a component of the γ-secretase complex^42–44^. It is plausible that RNA editing may influence splice-site choice in *KCNIP4*, lending to aberrant neuronal functioning through modulation of Kv4 channel functions and their subcellular location.

In contrast, higher levels of RNA editing were observed in genes that are essential for translation and mitochondrial protein translation. Several independent lines of evidence indicate mitochondrial dysfunction in schizophrenia, including mitochondrial hypoplasia, dysfunction of the oxidative phosphorylation system, and altered mitochondrial related gene expression^45–47^. Mitochondria dysfunction can severely affect neuronal activity, including synaptic connection, axon formation, and neuronal plasticity^46^. Among these genes are RNA Binding Motif Protein 8A (*RBM8A*), mitochondrial ribosomal protein 16 (*MRPS16*) and zinc finger protein 706 (*ZNF706)*. The molecular mechanisms of *RBMA* has been shown to control mRNA stability and splicing, translation and is located in the 1q21.1 copy-number variation associated with mental retardation, autism spectrum disorder, SCZ and microcephaly^48–50^. *MRPS16* was enriched for differentially edited sites located in its respective 3’UTR. Although our study focused on global patterns of A-to-I editing, a concerted approach with these human-specific sites, as they have been conducted in glutamatergic receptors, in the future will provide a more complete understanding of how RNA editing in these genes impact SCZ etiology.

We detected that edQTL are widespread in the brain and a substantial portion replicate between two brain regions. Approximately 30% of all RNA editing sites were associated with one or more nearby cis-regulatory variants. It is expected that the genomics of cis-edQTLs and their RNA editing sites align with context-specific regulation of editing, as indicated through overlap of edSNPs with regulatory elements, such as tissue-specific enhancers (**Fig. S17**) and mapping of RNA editing sites on genes, which predominately postnatally biased in neocortical gene expression (**Fig. S9**). Moreover, a fine mapping analysis was conducted between edQTL signatures and disease association in GWAS loci to identify RNA editing sites that may contribute to SCZ etiology. We identify six GWAS loci showing co-localization with edQTLs. Notably, genes *NGEF* and *ARL6IP4* replicated between brain regions. The edSNPs and editing sites for *NGEF* are located within 3’UTR regions and enhancer elements. *NGEF* is predominantly expressed in brain, particularly during early development, and shows substantial homology with the Dbl family, which have been shown to function as guanine nucleotide exchange factors for the Rho-type GTPases and are implicated in human cognitive function^51,52^. The editing site in *ARL6IP4* (also know as, splicing factor SRp25) causes a non-synonymous amino acid substitution (K/R) and affects a basic region in the protein that has not been ascribed a specific function. This lysine-to-arginine change does not alter the overall charge of the molecule, and represents a conservative change that may not affect the protein’s function substantially^53–54^. However, lysine residues can be sites of post-translational modification and thereby regulate protein function. We also identified GWAS-edQTL co-localization for *ENSA*, a gene which belongs to a highly conserved cAMP-regulated phosphoprotein family and is considered an endogenous regulator of ATP-sensitive potassium (K_ATP_) channels, which rest at the intersection of cell metabolism and membrane excitability^55^. K_ATP_ channels are open during states of low metabolic activity, resulting in hyperpolarization of the membrane, which has cytoprotective effects in neural tissues^55–56^. The diversity of KATP channel properties, resulting from differential molecular makeup, allows for exploitation by differential pharmacology, creating likely inroads towards new targeted pharmacological interventions.

In conclusion, our study reveals dynamic aspects of RNA editing in human brain tissue covering hundreds of SCZ cases and control samples, including two brain regions and two large primary cohorts used for discovery and validation. Strong reproducible evidence was identified for widespread dysregulation of RNA editing in SCZ, including under-editing of glutamate receptor activity and post-synaptic density genes, which show pyramidal neuronal cell type specificity as well as over-editing in genes involved in regulation of translation and translation initiation which are specific to interneuronal cell types. A significant fraction of these sites map to 3’UTR regions and enrich for common sequence motifs. Moreover, we characterize a large portion of RNA editing sites to be involved in cis-edQTLs in human brain tissue and further perform GWAS-edQTL colocalization analysis, which identified co-localization of 11 edQTLs with 6 GWAS loci. This result indicates that ~5% of all known SCZ risk loci co-localize with RNA editing, consistent with a causal role of RNA editing in risk for SCZ. While these results shed new light into the mechanisms underlying the neuropathophysiology of SCZ, additional molecular studies of aberrant RNA editing sites identified in the current study and their molecular mechanisms are required to fully appreciate their functional importance for SCZ neurobiology.

## MATERIALS AND METHODS

### Identification of RNA editing sites from human RNA-sequencing data

RNA-sequencing data generated from the human post-mortem ACC and DLPFC were obtained through the CommonMind Consortium (CMC). Additional RNA-sequencing data from human postmortem DLPFC were obtained through the NIH Human Brain Collection Core (HBCC). All fastq files were mapped to human reference genome hg19 using STAR version 2.4.0^1^ and the following parameters were optimized: chimSegmentMin=15; chimJunctionOverhangMin=15; outSAMstrandField=intronMotif. For each sample, this produced a coordinate-sorted BAM file of mapped paired end reads including those spanning splice junctions. Known RNA-edited sites were curated using the publicly available database, Rigorously Annotated Database of A-to-I RNA editing (RADAR)^2^. Nucleotide coordinates for these well documented editing sites were then used to extract reads from each sample using a customized perl script and the samtools mpileup function^3^. This approach quantifies the total amount of edited reads and the total amount of un-edited reads, which map to each RNA editing site in the RADAR database for each individual sample, thereby producing a rich source of editing information both within and across all samples. In order to identify a collection of high quality and high confidence sites, a series of thresholds were placed for each brain region and cohort, separately. The minimum base quality of 25, minimum mapping quality of 20 (that is, probability that a read is aligned to multiple locations), probability of misalignment = 0.01 (i.e., 99% probability that a read is correctly aligned in the genome), and minimum read coverage per edited site to be 20. The identification of RNA editing sites has previously been reported to be prone to these biases, therefore, it is likely that changing these parameters to be more lenient would increase the number of falsely predicted editing events. We also removed all known single nucleotide polymorphisms (SNPs) present in the SNP database (dbSNP; except SNPs of molecular type ‘cDNA’) and those within the 1000 Genomes Project. Finally, we required that an editing site must be present in at least 80% of all samples and subsequently, must have no more than 20% missing values per sample. The resulting RNA editing data frames for the CMC ACC and DLPFC samples contained 8.3% and 9.8% missing data respectively, and the data frame for HBCC DLPFC samples contained 7.6% missing data. All missing values were imputed using predictive mean matching method in the *mice* R package^4^, using five multiple imputations and 30 iterations. The resulting sets of sites identified from these RNA-sequencing data were subsequently referred to as *known* RNA editing sites and were used for downstream analysis.

### Identification of RNA editing sites from macaque RNA-sequencing data

To examine whether drug treatment effects were responsible for overall RNA editing levels observed in SCZ, we computed overall editing derived from an RNA-sequencing study of DLPFC tissue from Rhesus macaque monkeys. Antipsychotic administration, tissue dissection and RNA-sequencing data generation was previously described elsewhere^5^. In brief, subjects were randomly selected for four treatment groups: (1) high doses of haloperidol (4mg/kg/d), (2) low doses of haloperidol (0.14mg/kg/d), (3) clozapine (5.2mg/kg/d), (4) vehicle. Treatments were administered orally for six months. Following a six month treatment regime, monkeys were sacrificed using an overdose of barbiturate and transcardinally perfused with ice cold saline. DLPFC tissue was dissected from the dorsal and ventral banks of the principal sulcus (Area 46) and pulverized. Finally, gene expression data was generated using an identical RNA-sequencing protocol. Raw RNA-sequencing data was aligned to the macaque reference genome and transcriptome (mmul1) using STAR. Next, all well documented RNA editing sites in the RADAR database, which were annotated to the human reference hg19, were lifted over to the macaque reference mmul1 using the R library package rtracklayer^6^. These nucleotide coordinates were used to extract reads from each sample using the same customized perl script and the samtools mpileup function. We also carried out a series of matching thresholds in order to identify a collection of high confidence sites across all samples, as noted above. However, because so few sites were detected across all macaque and human samples, we restricted our comparative approach to measure the influence of medication on overall RNA editing levels, which is a threshold-free approach.

### Quantifying RNA editing levels

RNA editing levels were calculated for each sample, as previously described^7^. In brief, we define editing levels as the total number of edited reads at a specific RNA editing site (*i.e*., reads with G nucleotides) over the total number reads covering the site (*i.e*, reads with A and G nucleotides). The resulting metric is a continuous measure, ranging from 0 (*i.e*., a totally un-edited site) to 1 (*i.e*., a completely edited site). When computing *overall* RNA editing levels per sample, we did not impose any sequencing coverage criteria, but instead took all known sites from the RADAR database into account that were identified in each sample in our study to obtain the total amount of editing in each sample. In this way, overall RNA editing is defined as the total number of edited reads at all known RNA editing sites over the total number reads covering all sites for each sample. These measures were used to identify relationships between editing levels and SCZ and between editing levels and expression of editing enzymes.

### Differential RNA editing analysis

It is possible that RNA editing levels, similar to that observed in gene expression studies, are influenced by a number of biological and technical factors. By properly attributing multiple sources of RNA editing variation, it is possible to partially correct for some variables. Therefore, prior to differential RNA editing analysis, the editing variance for each site was partitioned into the variance attributable to each variable using a linear mixed model implemented in the R package variancePartition^8^. Under this framework, categorical variables (i.e., sample site, biological sex) are modeled as random effects and continuous variables (i.e., individual age, PMI) are modeled as fixed effects. Each site was considered separately and the results for all sites were aggregated afterwards. This approach enabled us to rationally include leading covariates into our downstream analysis, which may ultimately have an influence on differential RNA editing analysis. Subsequently, to identify sites with differential RNA editing levels between SCZ and control samples, we implemented linear model though the limma R package^9^ covarying for the possible influence of individual age, RNA integrity number (RIN), postmortem interval (PMI), sample site and sex. Significance values were adjusted for multiple testing using the Benjamini and Hochberg (BH) method to control the false discovery rate (FDR). Sites passing a multiple test corrected *P*-value < 0.05 were labeled significant.

### Supervised class prediction methods

In order to assess cross-validation of the SCZ-related sites, two prediction models were built using the differentially edited sites in the (1) DLPFC and (2) ACC derived from the CMC (here referred to as, training set) to predict case/control status (*i.e*. SCZ cases from control samples) from withheld DLPFC data derived from the HBCC (here referred to as, test set). Regularized regression models, including ElasticNet, Lasso and Ridge Regression were fit using the glmnet R package^10^. The penalty parameter lambda (λ) was estimated using 10-fold cross validation on each training set using the caret package in R, and ultimately set to lambda.min, the value of λ that yields minimum mean cross-validated error of the regression model. Once the models were fit, they were applied to RNA editing levels from the test set using the predict() function, which calculates the predicted log-odds of diagnostic status. Subsequently, area under the receiver operative curve (ROC) analysis was performed using the pROC package in R^11^. Classification accuracies were reported as area under the curve (AUC) on test samples to assess the precision of the models.

### Identification of enriched sequence motifs and RNA binding protein sites

Previous studies suggest that RNA editing events are mediated by RNA-binding proteins that recognize specific sequence motifs around the RNA editing sites. Therefore, we extracted ±20 bp long sequences relative to each differentially edited site in the ACC and DLPFC, both the discovery and validation cohorts, to discover potential motifs that may determine its interaction with the RNA editing enzyme complexes. These sequences were subjected to the Multiple Em Motif Elicitation (MEME) algorithm^12^ (http://meme-suite.org/). This method aims to detect motifs that are significant enriched within user defined list of sequences, regardless of their relative location to the editing sites. MEME was run using classic mode limiting the search to only the top 5 motifs whereby enrichment is measured relative to a (higher order) random model based on frequencies of the letters in the submitted sequences.

Sites enriched that shared a common sequence motif were then used to map binding sites of human RNA binding proteins (RBPs) using the RBPmap database^13^ (http://rbpmap.technion.ac.il/index.html). This produced a list of 37 motifs in the ACC and 36 motifs in the DLPFC discovery samples and 201 motifs in the DLPFC validation samples, which were independently submitted to RBPmap to identify motifs enriched in RBP targets from a database of 114 experimentally defined human motifs. The algorithm for mapping motifs on the RNA sequences is based on the Weighted-Rank approach, previously exploited in the SFmap web-server for mapping splicing factor binding, and was run in default mode.

### Genes enriched with differentially edited sites

In order to identify genes enriched with differentially edited sites, we corrected each gene for gene length. As gene length is strongly correlated with the number of detectable sites in each gene, we used a hyper-geometric test to examine over-representation of differentially edited sites within a particular gene while setting a rotating background to match the total number of detectable sites for each gene.

### Co-editing network analysis

To identify sites that are co-edited across SCZ and control samples, we applied unsupervised weighted gene co-expression network analysis (WGCNA)^14^. Signed networks were constructed for the CMC-derived ACC and DLPFC samples separately, and then again using HBCC-derived DLPFC samples, thus totaling three separate networks. To construct a network, the absolute values of Pearson correlation coefficients were calculated for all the possible editing site pairs and resulting values were transformed using a β-power of 8 for each network so that the final correlation matrix followed an approximate scale-free topology. The WGCNA cut-tree hybrid algorithm was used to detect sub-networks, or co-editing modules, within the global network with the following optimizations: minimum module size of 30 sites, tree-cut height of 0.999 and a deep-split option of 2. For each identified module, we ran singular value decomposition of each module’s editing matrix and used the resulting module eigengene (ME), equivalent to the first principal component, to represent the overall editing profiles for each module. Subsequently, modules with similar editing profiles were merged if ME values were highly correlated (R>0.9). Co-editing modules were interrogated for containing an over-representation of significantly differentially edited sites in SCZ using a Fisher’s Exact Test and an estimated odds-ratio in comparison to a background of all detected sites for each brain region and cohort. All pairwise tests were corrected using the BH method to control the FDR. To test whether ME values were significantly associated with SCZ, a linear model was applied covarying for individual age, RIN, PMI, sample site and sex using the *limma* package in R and all statistical tests were BH adjusted.

### Gene set and cell type enrichment analyses

All differentially edited sites passing a multiple test corrected *P*-value<0.05 and all co-editing network modules were subjected to functional annotation. The ToppFunn module of ToppGene Suite software^15^ (https://toppgene.cchmc.org/) was used to assess enrichment of GO ontology terms relevant to cellular components, molecular factors, biological processes and metabolic pathways using a one-tailed hyper-geometric distribution with a Bonferroni correction. A minimum of a three-gene overlap per gene set was necessary to be allowed for testing. Subsequently, modules were tested for over-representation of CNS cell type specific markers collected from a previously conducted single cell RNA-sequencing study^16^. In order for a gene to be labeled cell type specific, each marker required a minimum log_2_ expression of 1.4 units and a difference of 0.8 units above the next most abundance cell type measurement, as previously shown. Over-representation of cell type markers within co-editing modules was analyzed using a one-sided Fisher exact test to assess the statistical significance. All *P*-values, from all gene sets and modules, were adjusted for multiple testing using the BH procedure. We required an adjusted *P*-value <0.05 to claim that a cell type is enriched within a module.

### BrainSpan developmental gene set enrichment analysis

BrainSpan developmental RNA-seq data (www.brainspan.org) were summarized to GENCODE10 and gene-level RPKMs were used across 528 samples. From here, only the neocortical regions were used in our analysis -- dorsolateral prefrontal cortex (DFC), ventrolateral prefrontal cortex (VFC), medial prefrontal cortex (MFC), orbitofrontal cortex (OFC), primary motor cortex (M1C), primary somatosensory cortex (S1C), primary association cortex (A1C), inferior parietal cortex (IPC), superior temporal cortex (STC), inferior temporal cortex (ITC), and primary visual cortex (V1C). Samples with RIN <= 7 were filtered and removed from subsequent analysis. Genes were defined as expressed if they were present at an RPKM of 0.5 in 80% of the samples from at least one neocortical region at one major temporal epoch, resulting in 22,141 transcripts across 299 high-quality samples ranging from post-conception weeks (PCW) 8 to 40 years of age. Finally, expression values were log-transformed (log2[RPKM+1]).

Linear regression was performed at each of 22,141 transcripts, modeling gene expression as a continuous dependent variable, as a function of a binary ‘developmental stage’ variable. A total of 11 developmental stages were analyzed. A moderated *t*-test, computed using the limma R package, was used to determine which genes were uniquely over-expressed and under-expressed for each specific developmental stage against all other developmental stages. Models included gender, individual as a repeated measure and ethnicity as adjustment variables. Significance values were adjusted for multiple testing using the Benjamini and Hochberg (BH) method to control the false discovery rate (FDR). After the BH correction, genes with Q-value < 0.05 and an log_2_ fold change > 0.5 are defined as a genes highly expressed in a given developmental stage, whereas genes with Q-value < 0.05 and an log_2_ fold change < 0.5 are defined as a genes lowly expressed in a given developmental stage. These curated data formed the basis of our developmental stage gene set enrichment analysis. All processed data are available upon request. To test for overrepresentation of genes with differentially edited sites within a given gene set, a modified version of the GeneOverlap function in R was used so that all pairwise tests were multiple test corrected using the BH method. The Fisher’s exact test function also provides an estimated odds-ratio in comparison to a genome-wide background set to 27,546 transcripts.

### cis-edQTL analysis

A total of 11,242 high confidence sites in the ACC and 7,594 sites in the DLPFC edQTL (editing quantitative trait loci) were derived using genetically inferred Caucasian samples (ACC=368, DLPFC=426) across the 6.4 million genotyped and imputed markers with imputation score ≥ 0.8 and estimated minor allele frequency ≥ 0.05. For each of the RNA editing sites, we normalized editing levels by centering and scaling each measurement through subtracting out the mean editing level value and dividing by the standard deviation. Quantile normalization was then used to fit the distribution to a standard normal distribution. Subsequently, in order to map genome-wide edQTLs, we used a linear model on the imputed genotype dosages and standardized RNA editing levels using MatrixEQTL^17^. The RNA editing levels were covaried for sample site, sex, individual age, PMI, RIN and clinical diagnosis. In order to control for multiple tests, the FDR was estimated for all cis-edQTLs (defined as 100 KB between SNP marker and editing position), controlling for FDR across all chromosomes. We identified significant cis-edQTLs using a genome-wide significance threshold (q< 0.05). Max cis-edQTL (defined as the most significant eSNP per site, if any) were annotated for genomic regulatory elements according to ENCODE annotations implemented with the SNP nexus annotation tool (http://snp-nexus.org/index.html). To assess whether cis-edQTLs relate to known enhancer sequences, we tested for overlap between edQTLs and tissue-specific enhancer sequences from the FANTOM project covering 40 different tissues. We leveraged the SlideBase database^18^ (http://slidebase.binf.ku.dk/), which has well curated lists of enhancers found to be exclusively expressed across different tissues in humans. A permutation-based approach with 1,000 random permutations was used to determine statistical significance of the overlap between edSNP coordinates and enhancer regions using the R package regioneR^19^. A matching permutation analysis was used to assess edQTL overlap with previously generated expression QTL (eQTLs) in the ACC and DLPFC, which are publically available from synapse (syn7188631, syn7254151),

### GWAS-edQTL co-localization analysis

A total of 108 genome-wide significant (*P*<5.0×10^−8^) SCZ GWAS loci^20^, as defined by linkage disequilibrium r2 > 0.6 start and end positions, and edQTL sites overlapping those loci were considered for analysis. For those edQTL sites overlapping these GWAS loci, extended edQTL calling was performed using an increased window size in order to obtain edQTL statistics for the entire GWAS locus. GWAS and edQTL summary statistics (beta, standard error) for SNPs within each GWAS locus were used as input to coloc2^21^, and posterior probabilities for five hypotheses (H0, no GWAS or edQTL signal; H1, GWAS signal only; H2, edQTL signal only; H3, GWAS and edQTL signal but not co-localized; H4, co-localized GWAS and edQTL signals) were estimated for each locus. Loci with posterior probability for hypothesis H4 (PPH4) greater than 0.5 were considered to have co-localized GWAS and edQTL signals.

## Supporting information

## ACKNOWLEDGEMENTS

Data were generated as part of the CommonMind Consortium supported by funding from Takeda Pharmaceuticals Company Limited, F. Hoffman-La Roche Ltd and NIH grants R01MH085542, R01MH093725, P50MH066392, P50MH080405, R01MH097276, RO1-MH-075916, P50M096891, P50MH084053S1, R37MH057881,AG02219, AG05138, MH06692, R01MH110921, R01MH109677, R01 MH109897,U01 MH103392, and contract HHSN271201300031C through IRP NIMH. Brain tissue for the study was obtained from the following brain bank collections: the Mount Sinai NIH Brain and Tissue Repository, the University of Pennsylvania Alzheimer’s Disease Core Center, the University of Pittsburgh NeuroBioBank and Brain and Tissue Repositories, and the NIMH Human Brain Collection Core. CMC Leadership: Panos Roussos, Joseph D. Buxbaum, Andrew Chess, Schahram Akbarian, Vahram Haroutunian (Icahn School of Medicine at Mount Sinai), Bernie Devlin, David Lewis (University of Pittsburgh), Raquel Gur, Chang-Gyu Hahn (University of Pennsylvania), Enrico Domenici (University of Trento), Mette A. Peters, Solveig Sieberts (Sage Bionetworks), Thomas Lehner, Geetha Senthil, Stefano Marenco, Barbara K. Lipska (NIMH). DLPFC RNA-sequencing data, which formed the basis of the validation cohort, was provided by the National Institute of Mental Health Human Brain Collection Core (HBCC). Rhesus Macaque tissue was provided by Scott Hemby through the Stanley Medical Research Institute for Funding for Non-Human Primate Research; and funded by NIMH grant R01 MH074313.

## AUTHOR CONTRIBUTIONS

J.D.B., P.S., J.B.L. and M.S.B contributed to experimental design, study design and formulating the research question. J.D.B. and P.S. contributed the funding of this work. M.S.B. and A.D. contributed to data analysis. P.R., G.E.H., E.S., A.C, P.S., B.D. and J.D.B. contributed to leadership and supervision of various aspects of this work. M.S.B. and J.D.B. contributed to writing the manuscript and all authors contributed to completing the final version.

