## Supplementary material for "Dynamic landscape and genetic regulation of RNA editing in schizophrenia"

**AFFILIATIONS:** [1]Department of Psychiatry, [2]Department of Genetics and Genomic Sciences, [3]Seaver Autism Center for Research and Treatment, [4]Icahn Institute of Genomics and Multiscale Biology, [5]The Charles Bronfman Institute for Personalized Medicine, Icahn School of Medicine at Mount Sinai, New York, New York, 10029 USA; [6]Department of Genetics, Stanford University School of Medicine, Stanford, California, 94305, USA; [7]Pamela Sklar Division of Psychiatric Genomics, Icahn School of Medicine at Mount Sinai, New York, New York, 10029 USA; [8]Mental Illness Research, Education, and Clinical Center (VISN 2 South), James J. Peters VA Medical Center, Bronx, New York, 10468, USA; [9]Friedman Brain Institute, Icahn School of Medicine at Mount Sinai, New York, New York, 10029 USA; [10]Medical and Population Genetics, Broad Institute of MIT and Harvard, Cambridge, Massachusetts 02142 USA; [11]Department of Developmental and Regenerative Biology, Icahn School of Medicine at Mount Sinai, New York, New York, 10029 USA; [12]Department of Psychiatry, University of Pittsburgh School of Medicine, 3811 O'Hara Street, Pittsburgh, Pennsylvania 15213, USA; [13]Mindich Child Health and Development Institute, Icahn School of Medicine at Mount Sinai, New York, New York, 10029 USA.

**Supplemental Figures 1-18**, in brief:

**Figure S1.** Overview of the analytic pipeline.
**Figure S2.** Summary of patient statistics.
**Figure S3.** Overall RNA editing levels and ADAR expression in human samples.
**Figure S4.** Overall RNA editing levels and ADAR expression in macaque samples.
**Figure S5**. Characterization of detected RNA editing events.
**Figure S6.** Computing variance explained.
**Figure S7.** Relationship between RNA editing and gene expression.
**Figure S8.** Multivariate supervised classification.
**Figure S9.** BrainSpan developmental gene expression profiles.
**Figure S10.** Motif enrichment analysis.
**Figure S11.** Gene length versus RNA editing sites.
**Figure S12.** Module eigengene correlations with ADAR expression.
**Figure S13.** Validation of co-editing network analysis.
**Figure S14.** Quantile-Quantile plot restricting genetic variant search space within same gene.
**Figure S15.** Distance plots for edQTL analysis.
**Figure S16.** Correlations between RNA editing levels.
**Figure S17.** Tissue-specific enhancer enrichment analysis.
**Figure S18.** Cis-edQTLs that co-localize with GWAS loci.

**Figure S1**

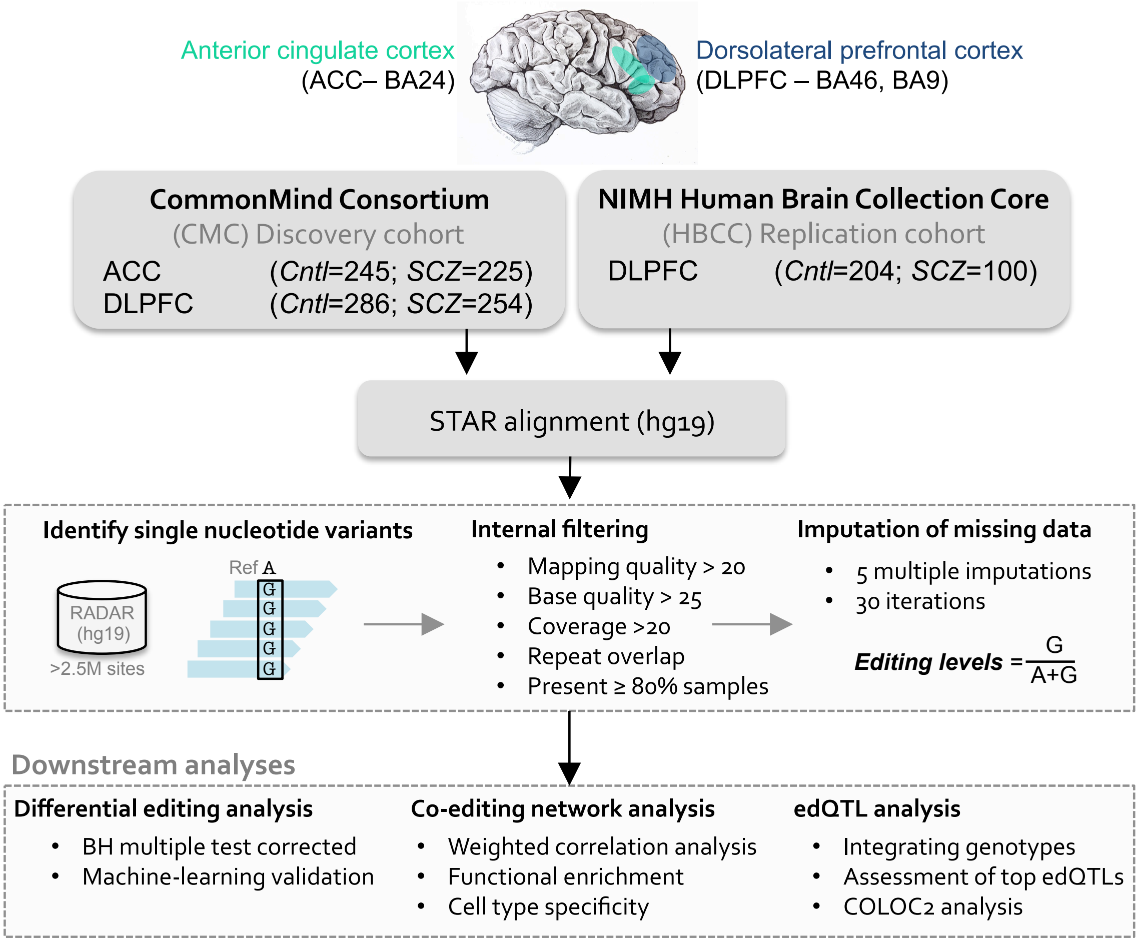


**Figure S1. Overview of the analytic pipeline used.** We leveraged RNA-sequencing data from post-mortem brain tissue collected and generated on behalf of the CommonMind Consortium (CMC). Two brain regions, including the ACC (SCZ=225, Controls=245) and the DLPFC (SCZ=254, Controls=286) were investigated, and together these samples served as the *discovery cohort*. In parallel, we also leveraged a completely separate, non-overlapping cohort consisting of post-mortem DLPFC tissue (SCZ=100, Controls=204) collected and generated on behalf of National Institute of Mental Health (NIMH) Human Brain Collection Core (HBCC). This second resource served as a *validation cohort* so as to cross-validate SCZ-related editing events identified from the CMC discovery samples. RNA-sequencing fastq files were aligned to the human reference genome and transcriptome (hg19) using STAR and bam files were sorted using samtools. We called RNA editing events from sorted bam files using the mpileup function in samtools together with customized perl scripts, which integrate all known RNA editing sites from the RADAR database. A series of internal filtering quality control metrics and imputations were computed before moving downstream to differential RNA editing, co-editing and edQTL analyses.

**Figure S2**

#### CommoMind Consoritum (CMC) Discovery cohort

| CMC DLPFC (*N* tot=540) | Control (N=286) | SCZ (N=254) | *P*-value |
|---|---|---|---|
| Site: *n* (MSSM/Penn/Pitt) | 162/38/82 | 141/58/55 | NA |
| Gender, female: *n* (%) | 124 (43%) | 94 (33%) | NA |
| Ethnicity, Caucasian: *n* (%) | 215 (75%) | 211 (74%) | NA |
| Age (years): | 65.86 ± 19.96 | 68.55 ± 16.97 | NA |
| PMI (hours): | 13.63 ± 8.16 | 20.5 ± 13.36 | 3.4e-08 |
| RIN: | 7.81 ± 0.87 | 7.40 ± 0.87 | 5.8e-06 |
| Brain weight (*g*): | 1244.99 ± 193.06 | 1231.41 ± 180.55 | NA |
| pH: | 6.57 ± 0.27 | 6.49 ± 0.27 | 0.02 |

| CMC ACC (*N* tot=470) | Control (*N*=245) | SCZ (*N*=225) | *P*-value |
|---|---|---|---|
| Site: *n* (MSSM/Penn/Pitt) | 143/24/78 | 125/41/59 | NA |
| Gender, female: *n* (%) | 100 (40%) | 81 (36%) | NA |
| Ethnicity, Caucasian: *n* (%) | 184 (75%) | 184 (81%) | NA |
| Age (years): | 64.94 ± 20.42 | 67.61 ± 17.24 | NA |
| PMI (hours): | 13.59 ± 7.87 | 19.85 ± 12.50 | 1.5e-06 |
| RIN: | 7.58 ± 0.85 | 7.34 ± 0.76 | 0.001 |
| Brain weight (*g*): | 1246.09 ± 193.72 | 1241.07 ± 174.17 | NA |
| pH: | 6.59 ± 0.26 | 6.51 ± 0.26 | 0.03 |

#### NIMH Human Brain Collection Core (HBCC) Replication cohort

| HBCC DLPFC (*N* tot=317) | Control (*N*=217) | SCZ (*N*=100) | *P*-value |
|---|---|---|---|
| Gender, female: *n* (%) | 60 (27%) | 35 (35%) | NA |
| Ethnicity, Caucasian: *n* (%) | 90 (41%) | 38 (38%) | NA |
| Age (years): | 30.05 ± 19.99 | 49.92 ± 13.34 | 9.2e0-7 |
| PMI (hours): | 29.04 ± 13.34 | 36.04 ± 22.00 | 0.005 |
| RIN: | 7.66 ± 0.89 | 7.32 ± 0.89 | 0.002 |
| Brain weight (*g*): | 1351.31± 219.53 | 1348.43 ± 162.12 | NA |
| pH: | 6.49± 0.29 | 6.37 ± 0.22 | 0.007 |

**Figure S2. Summary of patient statistics.** Patient level covariates were recorded and compared between SCZ cases and control samples, separately for each brain region (ACC and DLPFC) and cohort (discovery and validation). A Shapiro-Wilk test was used to assess normality of covariates and either a Student's *t*-test or Mann-Whitney U-test was implemented accordingly.

**Figure S3**

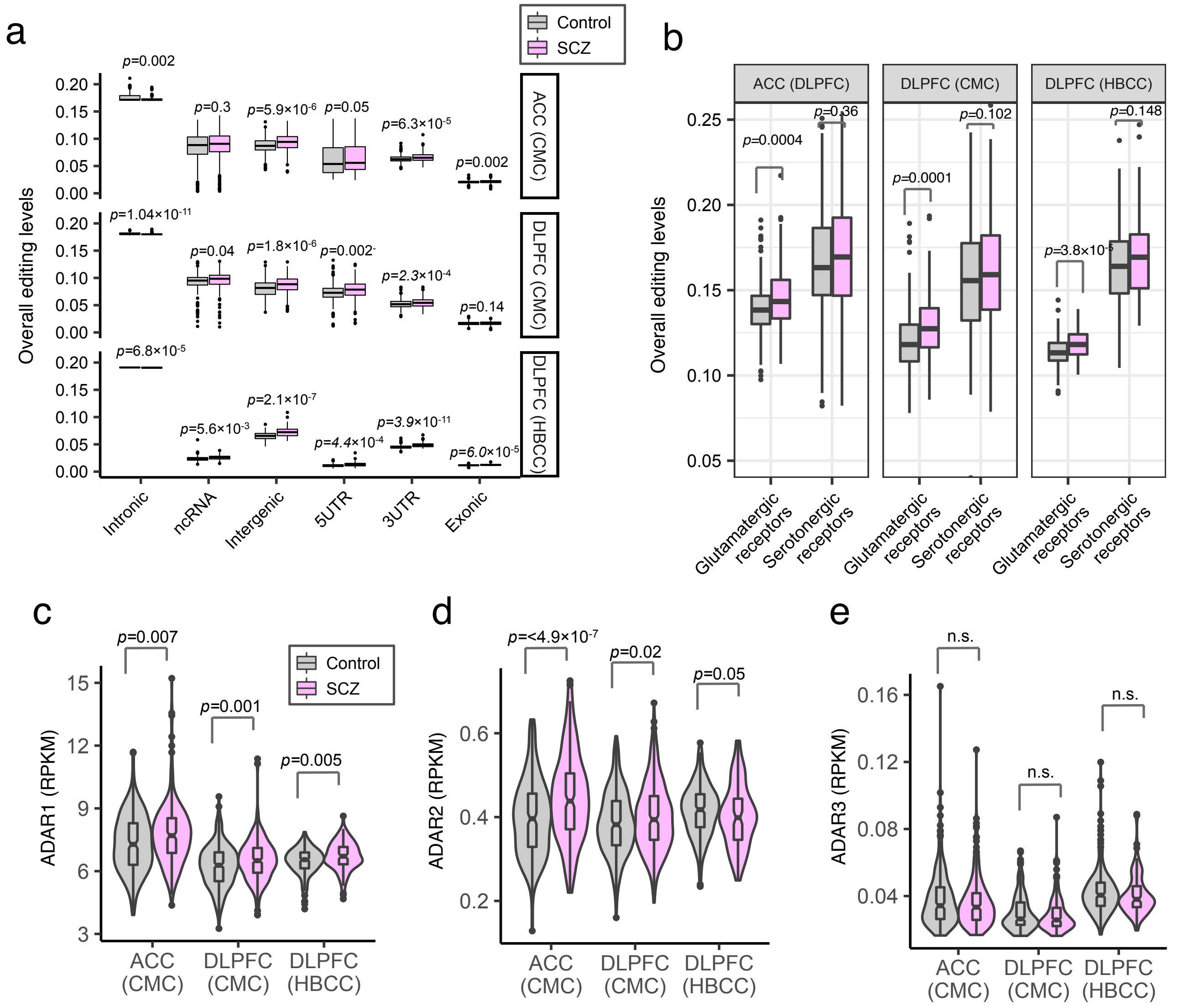


**Figure S3. Overall RNA editing and ADAR expression in human samples.** Overall RNA editing levels are computed separately for (**a**) each discrete genic region and (**b**) a priori defined glutamatergic and serotonergic receptor activity gene sets (GO:009589 and GO:0008066, respectively). Reads per kilobase of transcript per million mapped reads (RPKM) expression levels for (**c**) *ADAR1*, (**d**) *ADAR2* and (**e**) *ADAR3*. For all comparisons, a Mann-Whitney U test was used to test significance between groups. Note that *ADAR2* expression for HBCC samples is trending opposite to that in CMC samples.

**Figure S4**

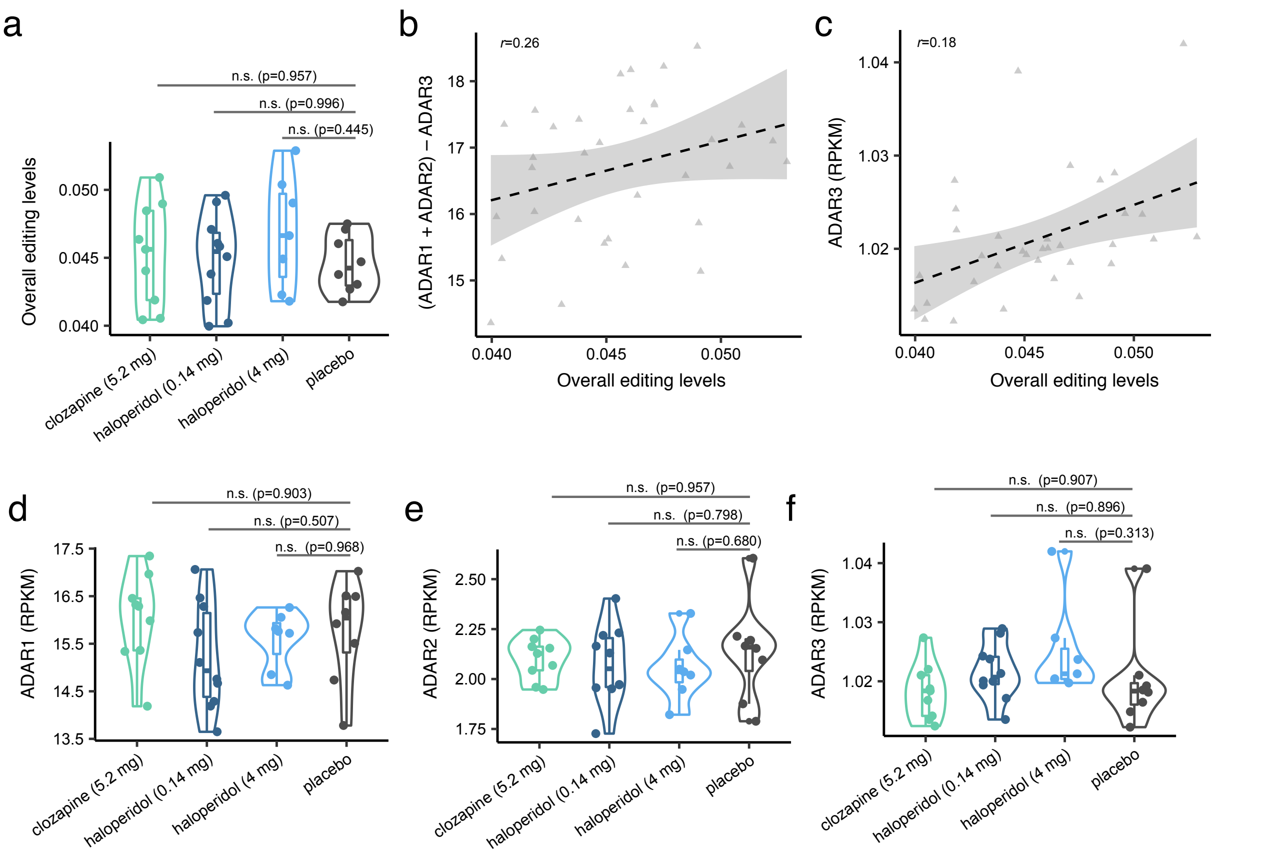


**Figure S4. Overall RNA editing and ADAR expression in macaque samples. (a)** Overall RNA editing levels are computed separately for DLPFC samples treated with different antipsychotic medications and dosages. Variance of overall RNA editing levels explained by (**b**) ADAR1 and ADAR2 and (**c**) ADAR3 Reads per kilobase of transcript per million mapped reads (RPKM) expression levels. RPKM expression levels for (**d**) ADAR1, (**e**) ADAR2 and (**f**) ADAR3. For all comparisons, a Dunnett's multiple comparison of means test was used comparing each treatment group relative to placebo.

#### Figure S5

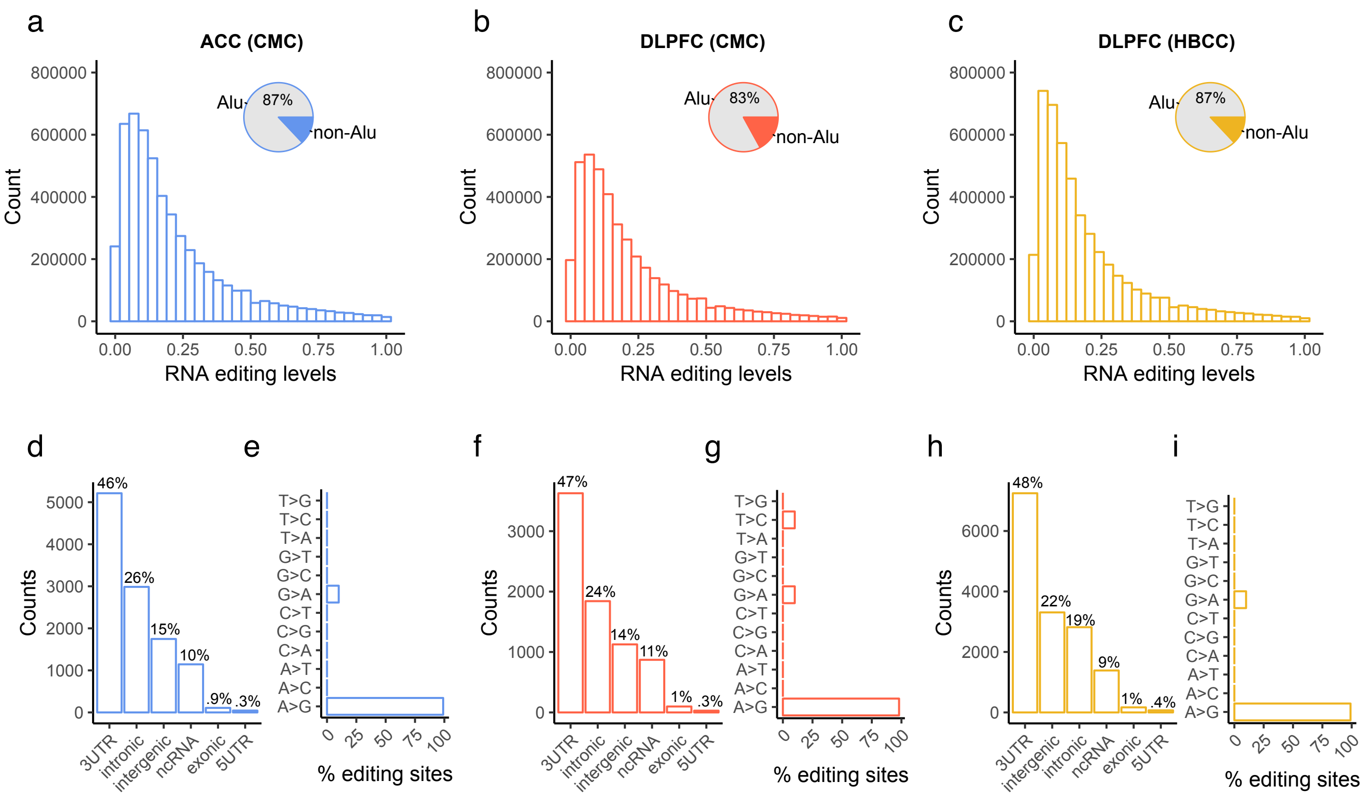


**Figure S5. Characterization of detected RNA editing events.** Frequency distributions of RNA editing levels for all detected sites in the (**a**) ACC and (**b**) DLPFC CMC samples and (**c**) DLPFC HBCC samples. Inset pie charts indicate the total fraction of all detected sites that map to Alu repeat elements. The total number of RNA editing events were summarized within each genic region and nucleotide conversion rates were assessed for the (**d-e**) ACC and (**f-g**) DLPFC CMC samples and (**h-i**) DLPFC HBCC samples.

**Figure S6**

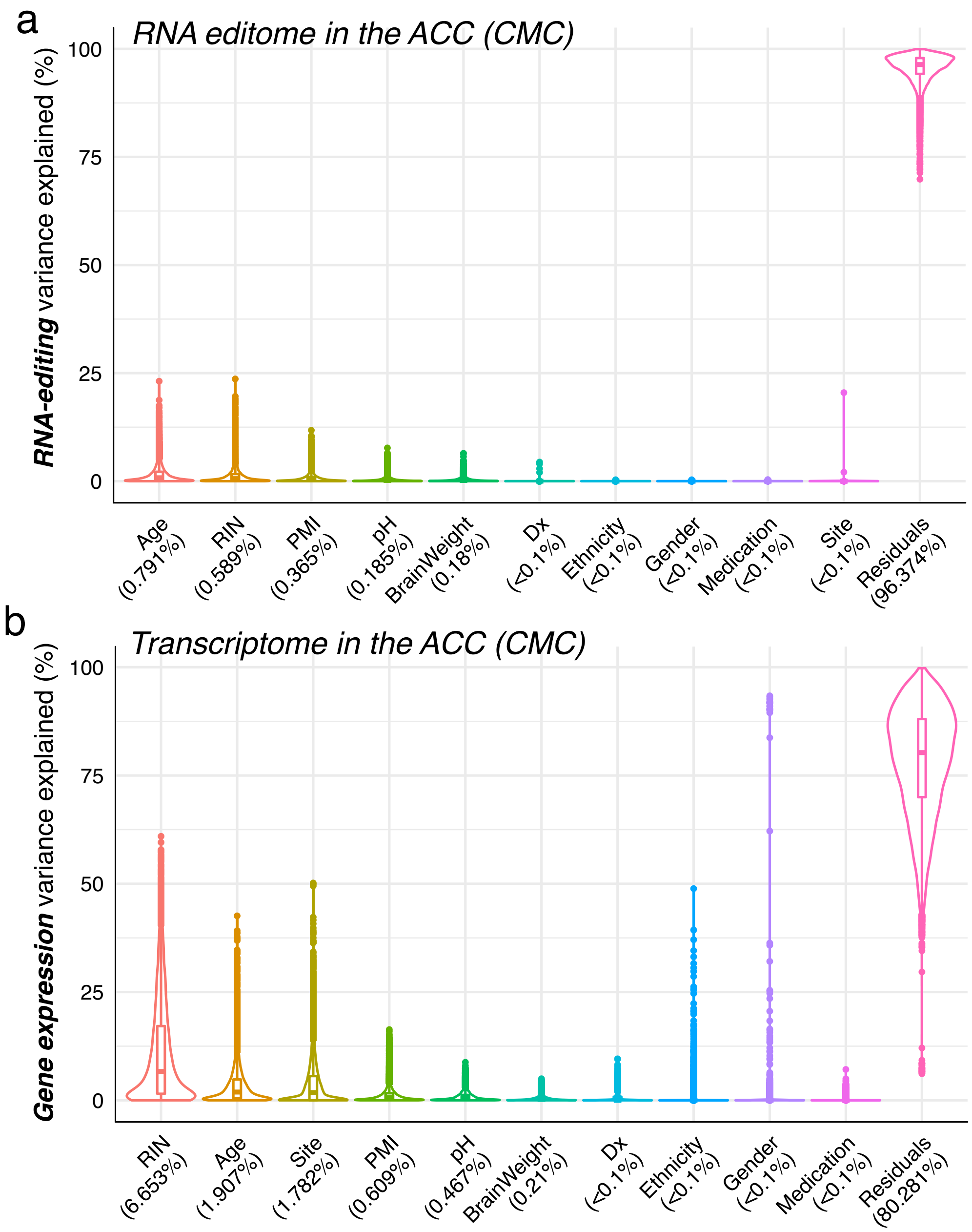


**Figure S6. Computing variance explained.** Variance explained according to nine covariates, which represent potential technical, biological and clinical sources of variability. We used the linear mixed model framework of the varianceParition R package to quantify variability explained in the (**a**) RNA editome and (**b**) transcriptome across all ACC samples. The dynamic range of transcriptome data finds stronger relationships and is under greater influence with the recorded covariate than RNA-editing measurements. Results from the CMC DLPFC samples and HBCC DLPFC samples were highly similar to those depicted here for the ACC and are not shown.

#### Figure S7

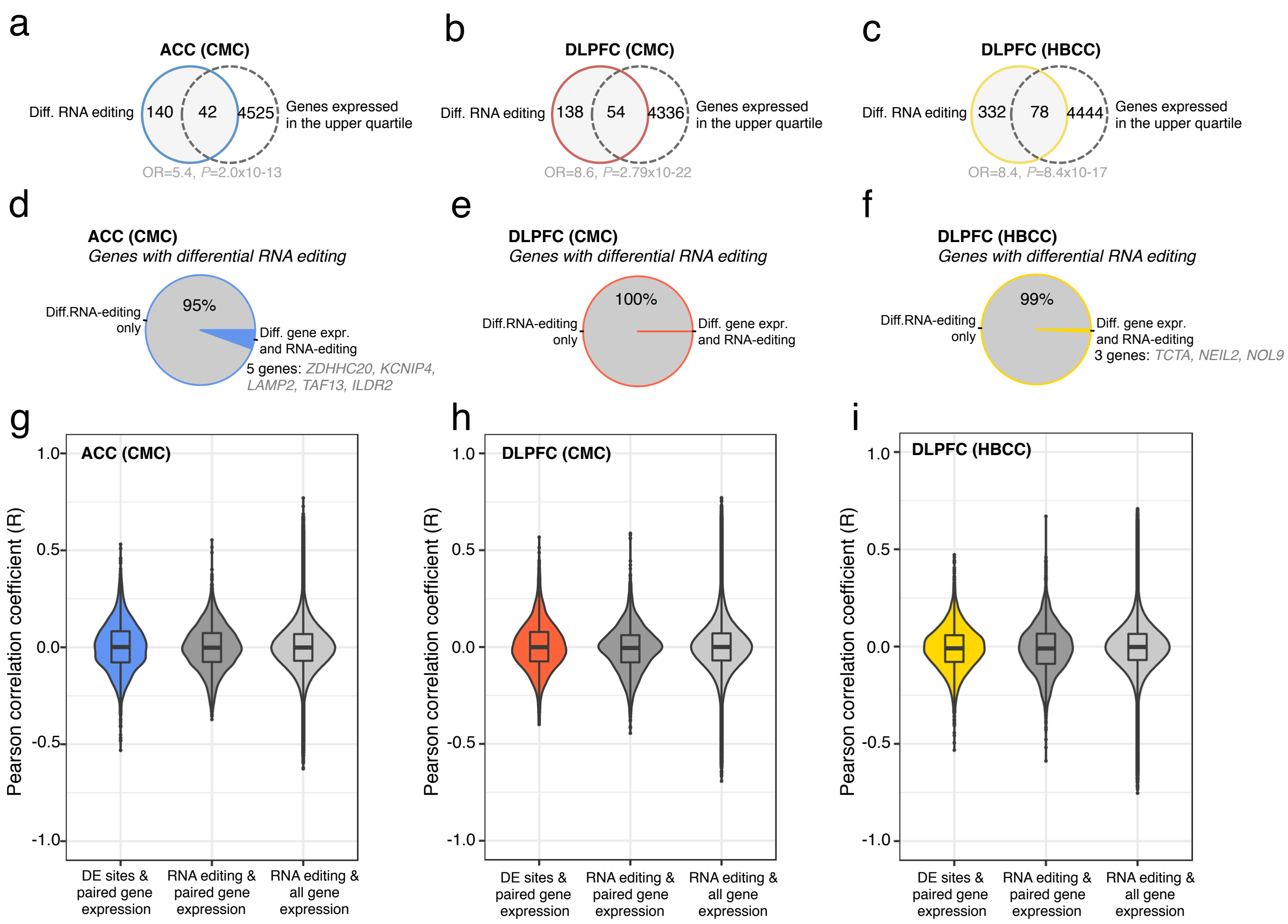


**Figure S7. Relationship between RNA editing and gene expression.** Enrichment analysis assessing whether genes harboring differentially edited sites overlap with genes that are highly expressed in the (**a**) ACC and (**b**) DLPFC CMC samples and (**c**) DLPFC HBCC samples. Genes were labeled 'highly expressed' if they had an average gene expression value across all samples that was in the upper $3^{rd}$ quartile. A Fisher's exact test was used to compute overlap significance and estimated odds-ratios. Overlap analysis assessed whether genes with differential RNA editing sites also displayed differential expression in the (**d**) ACC and (**e**) DLPFC CMC samples and (**f**) DLPFC HBCC samples. Inset gene symbols indicate genes which harbor differential editing sites and are dysregulated in SCZ cases compared to controls. Correlation analysis of RNA editing levels were compared to gene expression levels in three instances: 1) for differentially RNA edited sites relative to their respective gene expression levels; 2) all RNA editing sites relative to their respective gene expression levels; 3) all RNA editing sites relative to gene expression levels other than their respective gene. This analysis was carried out for the (**g**) ACC and (**g**) DLPFC CMC samples and (**i**) DLPFC HBCC samples.

#### Figure S8

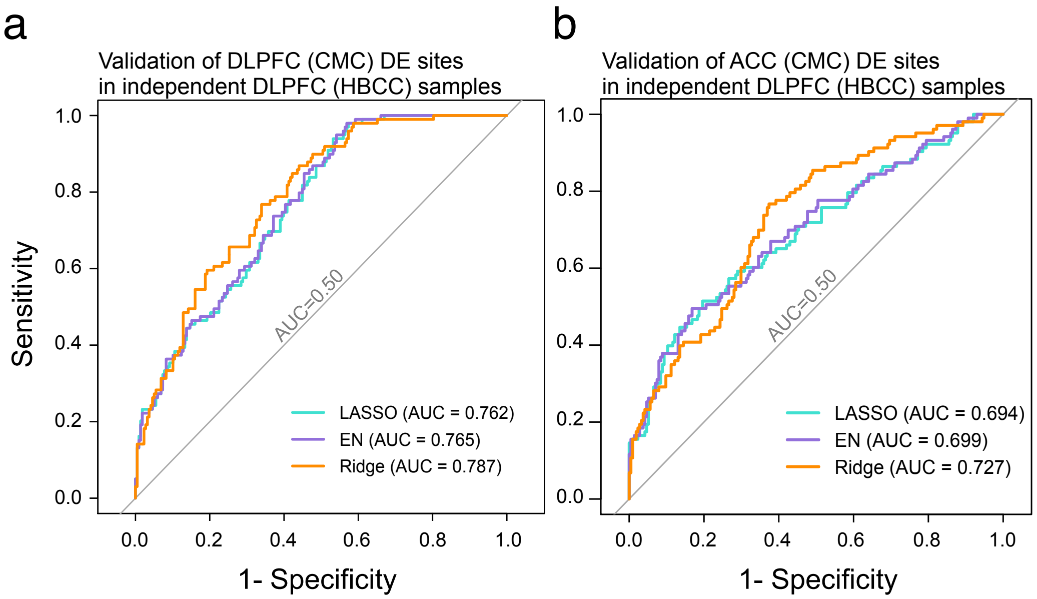


**Figure S8. Multivariate supervised classification.** Three regularized regression techniques, including ElasticNet (EN), Lasso and Ridge Regression were fit using the glmnet R package in order to assess cross-validation of the schizophrenia (SCZ)-related sites derived from CMC samples in withheld HBCC samples. Two prediction models were built using the differentially edited sites in the (**a**) DLPFC and (**b**) ACC derived from the CMC training sets to predict case/control status from withheld DLPFC data derived from the HBCC test set. Area under the receiver operator curve (AUC) values are used to assess the overall precision of these models. Ridge Regression achieved 78% and 72% prediction accuracy when using altered RNA editing and samples derived from DLPFC and ACC training data, respectively, to predict DLPFC HBCC test set samples.

#### Figure S9

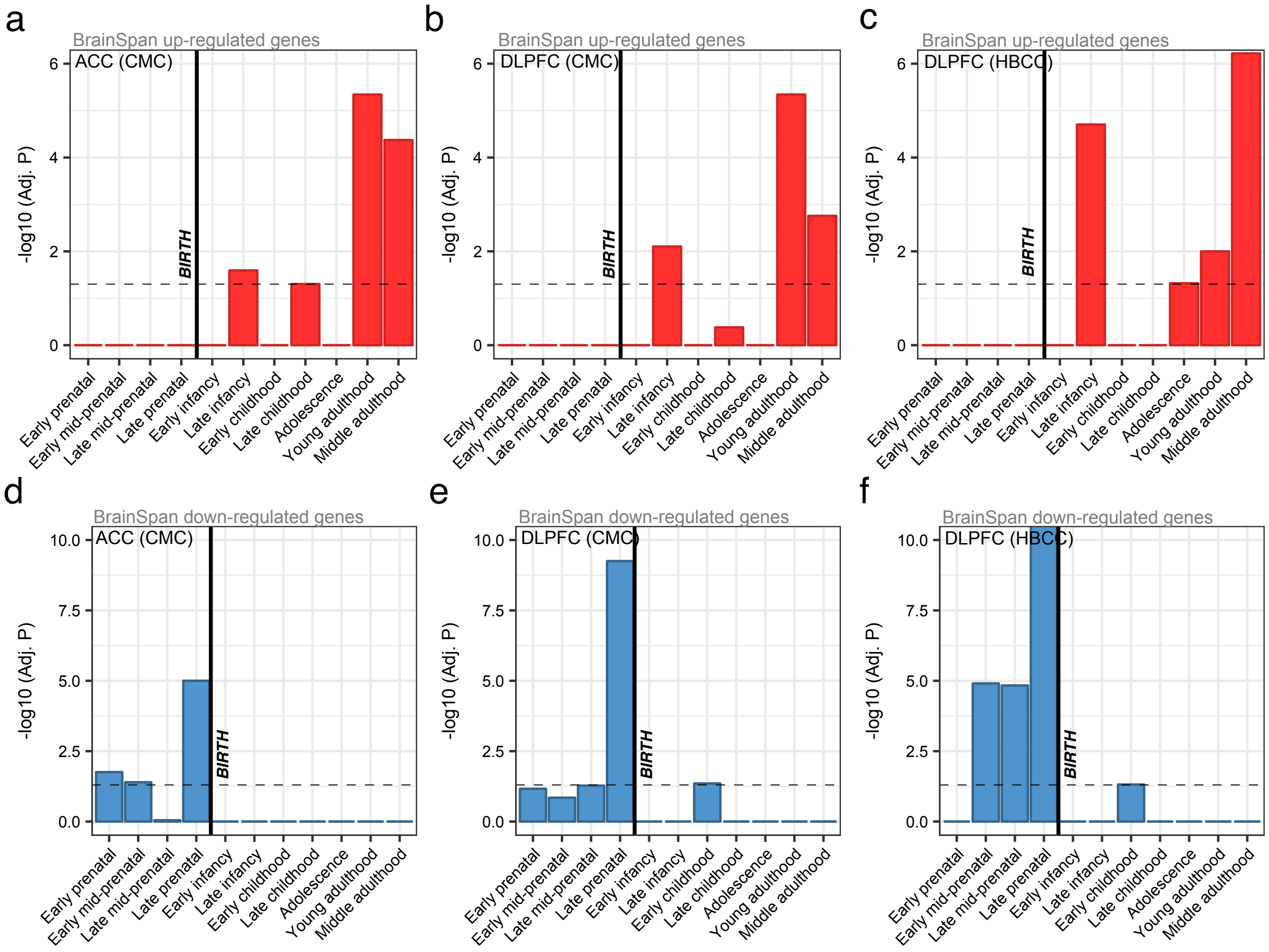


**Figure S9. BrainSpan developmental gene expression profiles.** Enrichment analysis examined whether differentially edited sites in schizophrenia (SCZ) mapped to genes with specific developmental trajectories. A total of 11 developmental stages (x-axis) were analyzed and gene sets, indicating whether genes are over-expressed or under-expressed at each stage relative to all other stages, were used to compute overlap and enrichment. A Fisher's exact test was used to calculate significance for all tests (y-axis). A consistent enrichment of differentially edited sites mapping to genes, which are highly expressed during young and middle adulthood was observed for the (**a**) ACC and (**b**) DLPFC CMC samples and (**c**) DLPFC HBCC samples. Additionally, we observed that these genes were also predominately under-expressed during the fetal period for the (**d**) ACC and (**e**) DLPFC CMC samples and (**f**) DLPFC HBCC samples, indicating that the developmental expression properties of these genes gradually increase in expression from early prenatal periods and peak during adulthood.

**Figure S10**

a

b

c

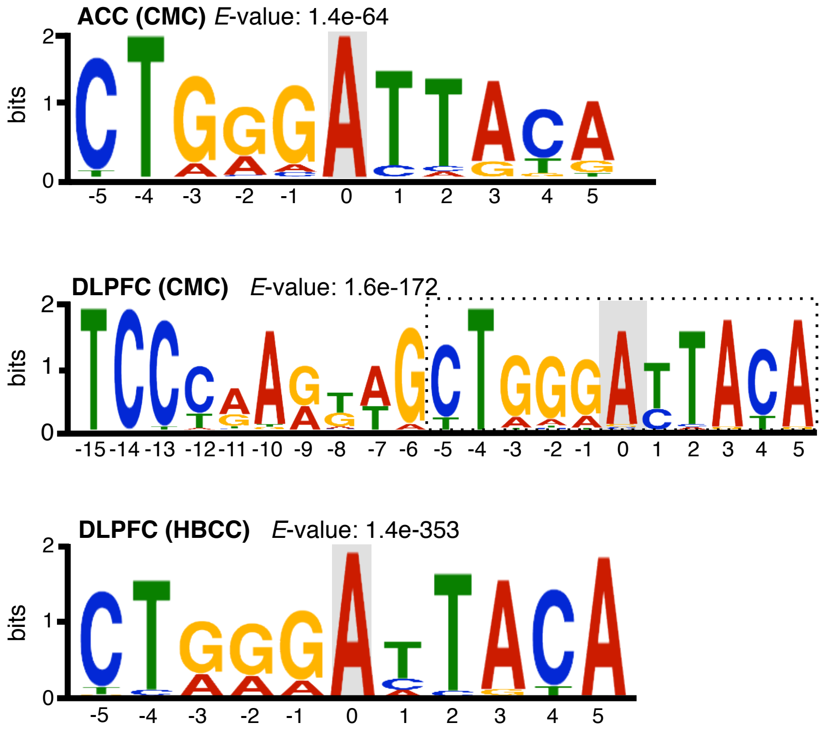


**Figure S10. Motif enrichment analysis.** All differentially edited sites were assessed for motif enrichment analysis ± 20bp from the editing site using MEME. Consistent and strong enrichment was observed for 10bp motif (± 5bp from the editing site) for differentially edited sites derived from the (**a**) ACC and (**b**) DLPFC CMC samples and (**c**) DLPFC HBCC samples. The overall height of each stack indicates the sequence conservation at that position (measured in bits), whereas the height of symbols within the stack reflects the relative frequency of the corresponding nucleic acid at that position. The editing site is outlined in grey.

#### Figure S11

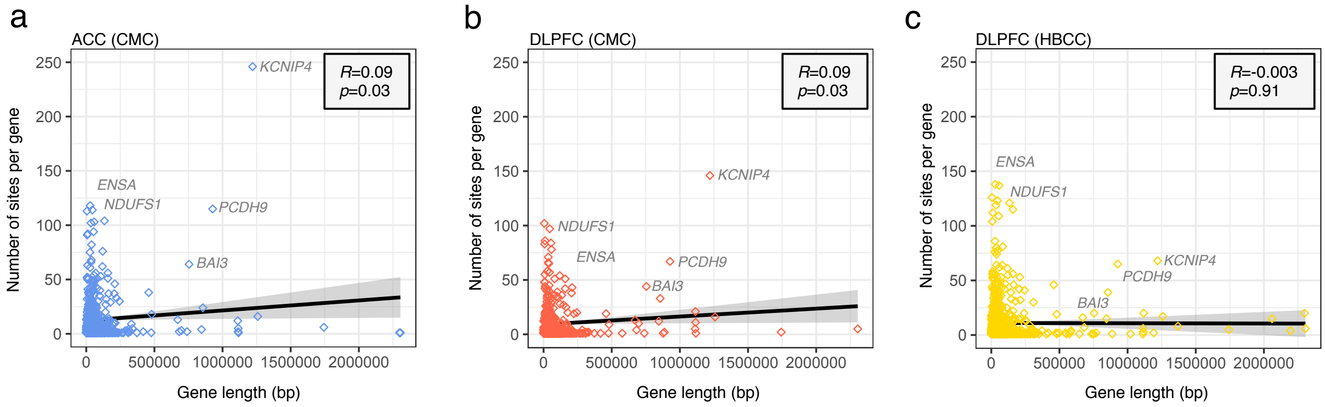


**Figure S11. Gene length versus RNA editing sites.** The total number of detected RNA editing events correlates with gene length for sites identified in the (**a**) ACC and (**b**) DLPFC CMC samples and (**c**) DLPFC HBCC samples.

#### Figure S12

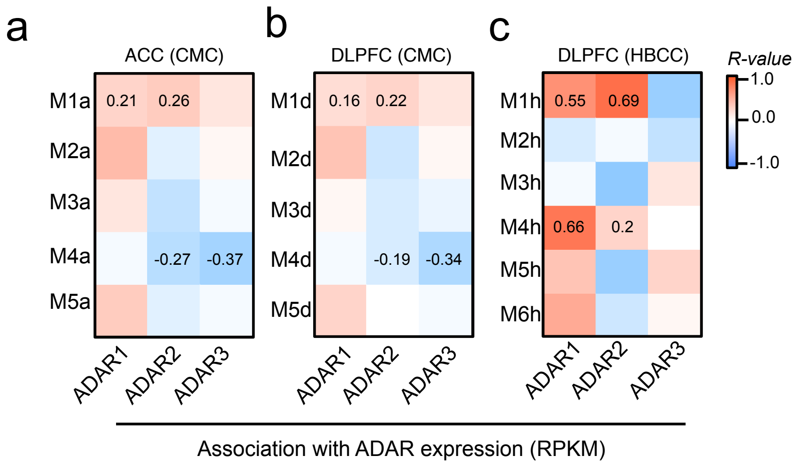


**Figure S12. Module eigengene correlations with ADAR expression.** Pearson correlation coefficients between module eigengene values and ADAR expression values for co-editing networks identified in the the (**a**) ACC and (**b**) DLPFC CMC samples and (**c**) DLPFC HBCC samples. Expression is quantified as the number of RNA-seq reads per kilobase of transcript per million mapped reads (RPKM). Correlation coefficients for modules of interest are presented in each corresponding cell.

### Figure S13

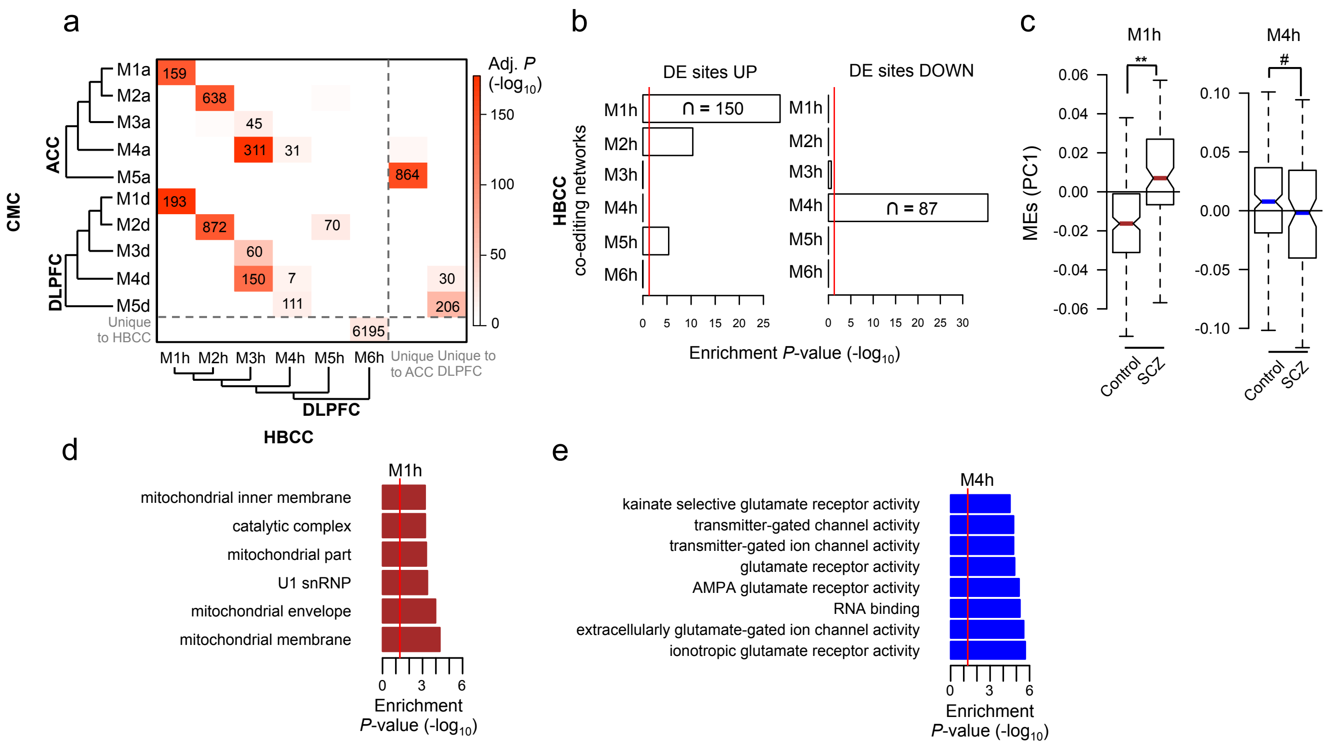


**Figure S13. Validation of co-editing network analysis.** Unsupervised co-editing network analysis was applied to DLPFC HBCC samples (validation samples). **(a)** Overlap analysis of co-editing modules derived from these validation samples were compared to those previously identified within the ACC and DLPFC discovery. Unsupervised clustering was used to group modules by module eigengene (ME) values using Pearson's correlation coefficient and Ward's distance method. Significance of overlap was computed using a Fisher exact test corrected for multiple comparisons. (**b**) Enrichment analysis of differentially edited sites within co-editing networks in the DLPFC HBCC samples. (**c**) Assessment of ME values for modules M1h (over-edited) and M4h (under-edited). Differential ME analysis was conducted using a linear model and covarying for age, RIN, PMI, sample site and gender. (**d**) The top functional enrichment terms for the (**e**) over-edited module M1h and (e) under-edited module M4h.

**Figure S14**

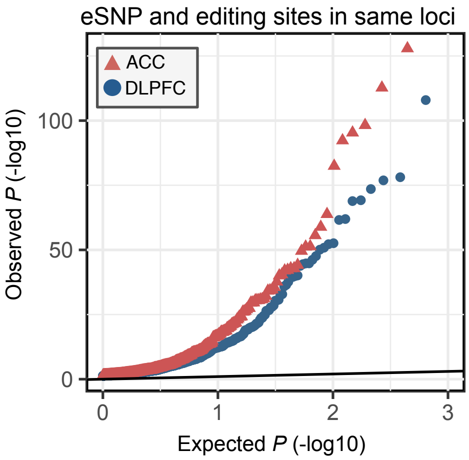


**Figure S14. Quantile-Quantile plot.** Quantile-quantile plot for association testing *P* values between RNA editing sites and genetic variants in the same gene as each editing site for the ACC (red) and DLPFC (blue).

### Figure S15

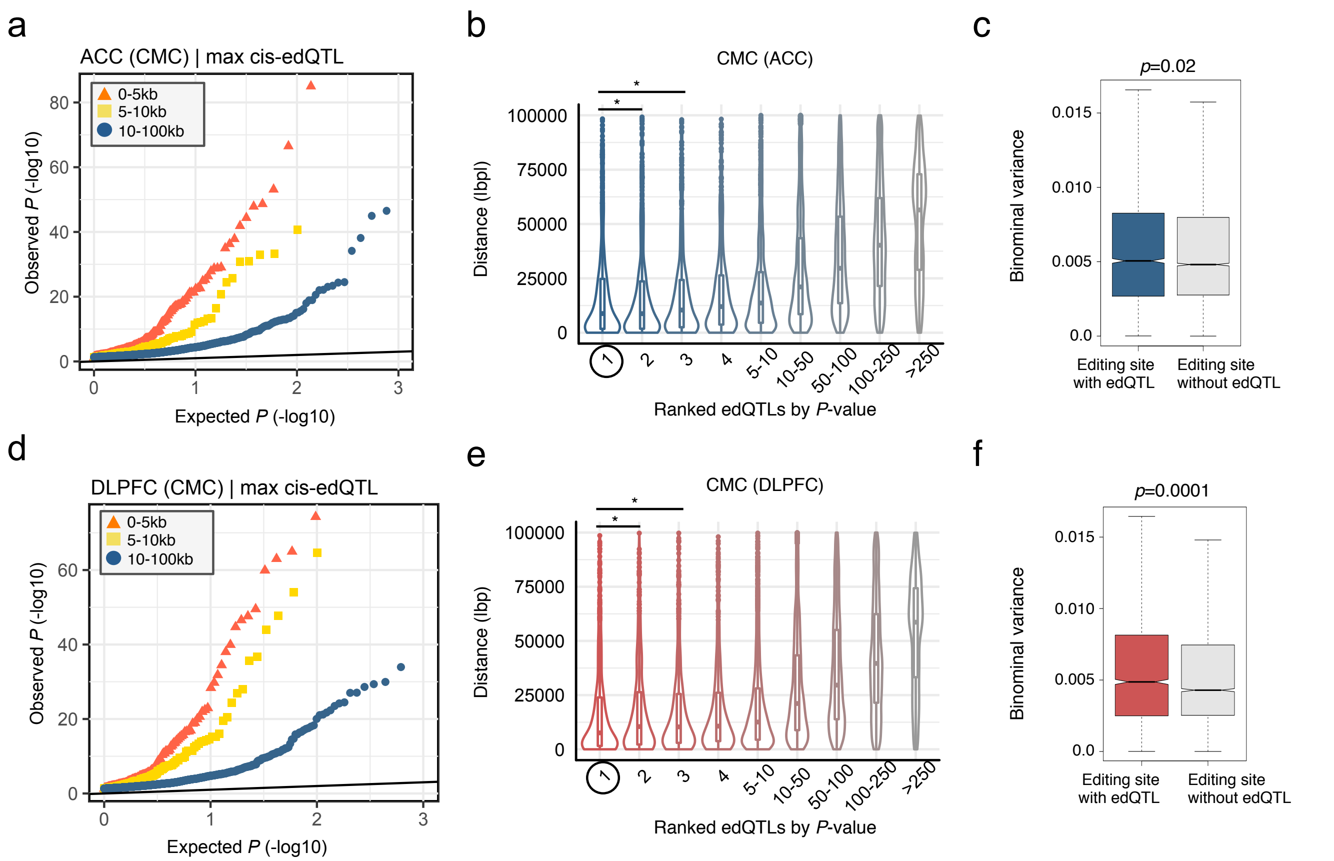


**Figure S15. Distance plots for edQTL analysis.** Quantile-quantile plot for association tests between edQTLs and additional editing sites that fell within 5 kb (orange), between 5 kb and 10 kb (gold), and between 10 kb and 100 kb (blue) from the original best-associated editing site for (**a**) ACC and (**d**) DLPFC samples. Violinplots quantify significant differences in proximity of each edSNP relative to its corresponding editing site for each max-edQTL relative to all other edQTLs for that same editing site in the (**b**) ACC and (**e**) DLPFC. edQTLs are ranked by significance, with the max-edQTL prioritized as number one and the second most significant edQTL prioritized as number two, and so on. Furthermore, binomial variance analysis indicates RNA editing sites with edQTLs display more variances than those which do not have edQTLs in the (**c**) ACC and (**f**) DLPFC. For all comparisons, a Mann-Whitney U test was used to test significance between groups.

#### Figure S16

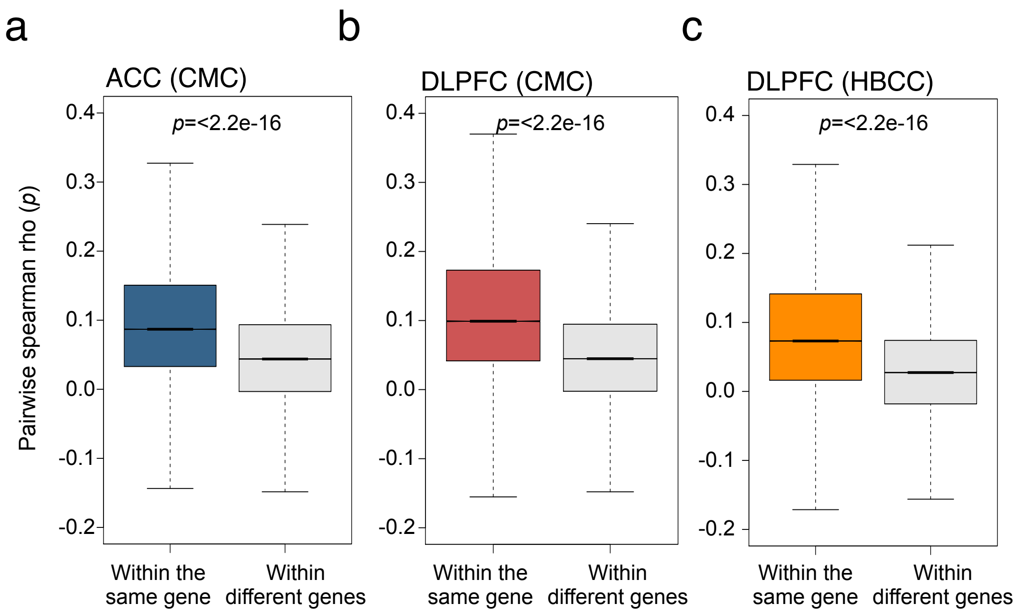


**Figure S16. Correlations between RNA editing levels.** Spearman correlation coefficient computed a series of pairwise associations between RNA editing levels for sites within the same gene as well as pairwise associations between RNA editing levels for sites in all other genes. Higher correlations were observed between sites in the same gene, in the (**a**) ACC and (**b**) DLPFC CMC samples and (**c**) DLPFC HBCC samples.

**Figure S17**

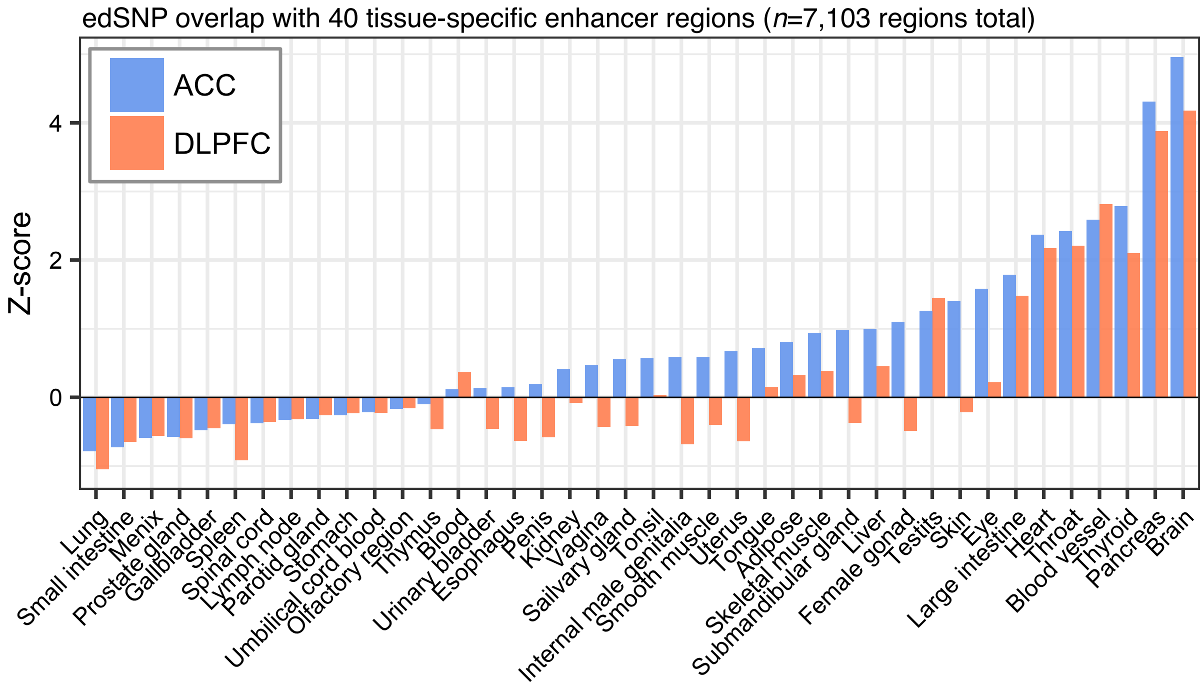


**Figure S17. Tissue-specific enhancer enrichment analysis.** Genomic coordinates for edSNPs were overlapped with tissue-specific enhancer regions derived from 40 different human tissues; data from the FANTOM project. The regioneR R package was used test overlaps of genomic regions based on permutation sampling. We repeatedly sampled random regions from the genome 1000 times, matching size and chromosomal distribution of the region set under study. By recomputing the overlap with the enhancer features in each permutation, statistical significance of the observed overlap was computed. We observed enrichment for many tissues, but the strongest enrichment was for brain tissue in the (**a**) ACC and (**b**) DLPFC.

#### Figure S18

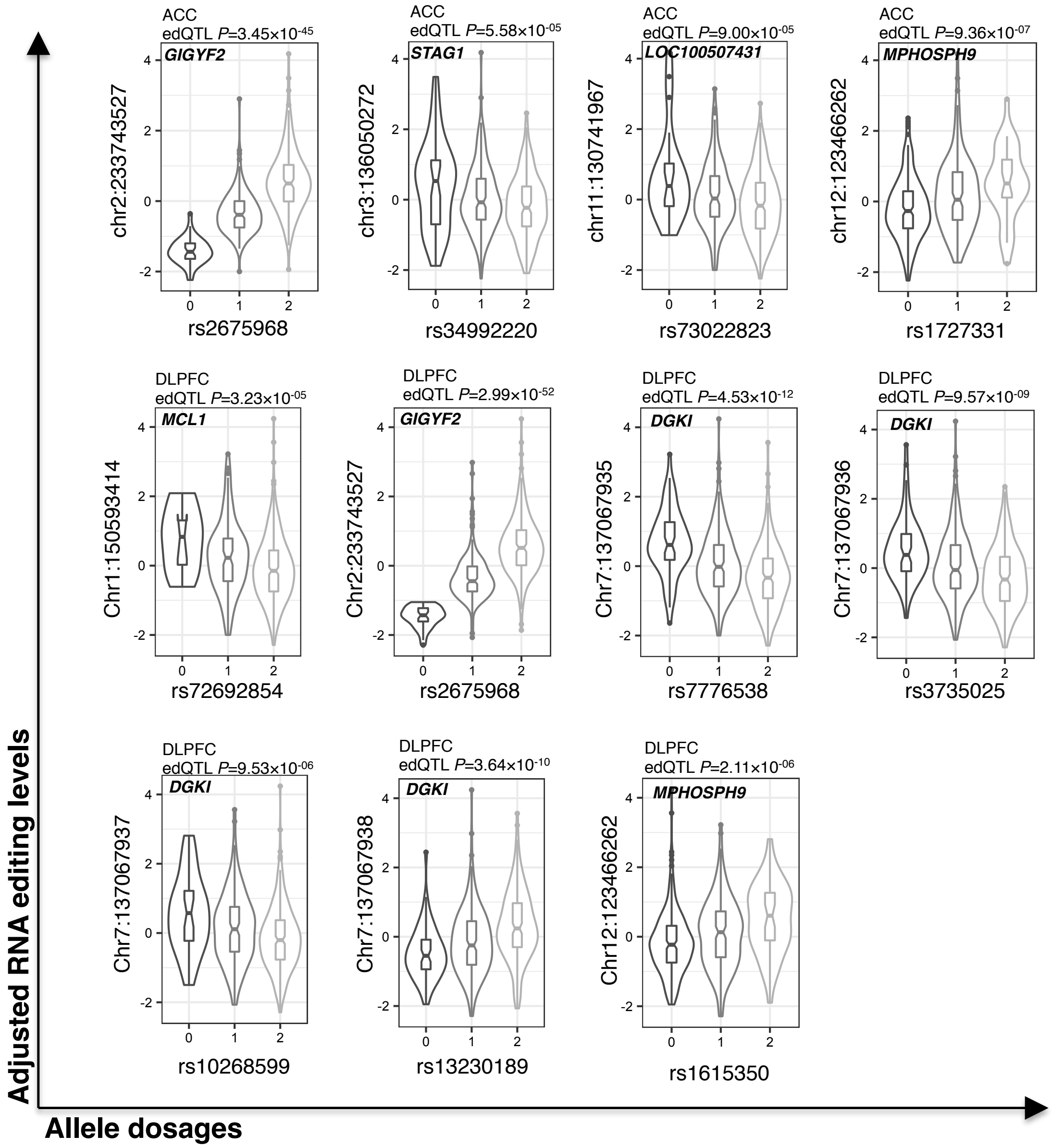


**Figure S18. Cis-edQTLs that co-localize with GWAS loci.** Cis-edQTL plots for GWAS-edQTL co-localized loci. The allelic effect of the SNPs on editing levels are shown by boxplots within violin plots. Violin plot shows the density plot of the data on each side, the lower and upper border of the box correspond to the first and third quartiles, respectively, the central line depicts the median, and whiskers extends from the borders to ±1.5xIQR, where IQR stands for inter-quantile range, the distance between the first and third quantiles.
